# Molecular Jackhammers Eradicate Cancer Cells by Vibronic-Driven Action

**DOI:** 10.1101/2023.01.25.525400

**Authors:** Ciceron Ayala-Orozco, Diego Galvez-Aranda, Arnoldo Corona, Jorge M. Seminario, Roberto Rangel, Jeffrey N. Myers, James M. Tour

## Abstract

Through the actuation of vibronic modes in cell-membrane-associated aminocyanines, a new type of molecular mechanical action can be exploited to rapidly kill cells by necrosis. This is done using near-infrared light, a low energy source hitherto thought to be insufficient to permit molecular mechanical disruption of a cell membrane. Vibronic-driven action (VDA) is distinct from both photodynamic therapy and photothermal therapy in that the VDA mechanical effect on the cell membrane is not retarded by high doses of inhibitors of reactive oxygen species (ROS), and VDA does not itself induce an increase in the temperature of the media; it is also unaffected by cooling the media to 2 °C. The picosecond concerted whole-molecule-vibrations of VDA-induced mechanical disruption can be done with very low concentrations (500 nM) of the aminocyanines or low doses of light (12 Jcm^-2^, 80 mWcm^-2^ for 2.5 min) to cause *in vitro* necrotic cell death in >99% of human melanoma cells. The effect is also studied *in vivo* in murine B16-F10 and human A375 melanoma in mice, underscoring the high efficiency of this approach, achieving a survival rate of 60% at day 120, and 50% of the mice becoming tumor free. The molecules that destroy cell membranes through VDA are termed molecular jackhammers (MJH) because they undergo concerted whole-molecule vibrations. Different than traditional chemotherapy, it is unlikely that a cell could develop a resistance to molecular mechanical forces, thereby providing a new modality for inducing cancer cell death.

## Introduction

We previously used Feringa-type unidirectional rotating molecular motors that bear a rotor and stator that can be activated by ultraviolet (UV) or visible light (Vis) to open pores in cell membranes resulting in rapid necrotic death.^1^ Upon the 2-3 MHz unidirectional rotations in these molecular motors, drilling action on cellular membranes develop holes in the cell, then blebbing and death within seconds to minutes of activation. This necrotic action is unaffected by inhibitors of radical oxygen species (ROS).^2^ Slower versions of the Feringa motors, where the rotation rates are on the order of 1-100 Hz, do not show such cell-killing action, further ruling out a thermal effect in the killing.^2^ We have more recently studied visible-light-activated Dube-hemithioindigo switches and motors that operate in the kHz regime,^3^ and although still these are too slow to mechanically kill cells in our studies, they can induce ROS resulting in slower apoptotic cellular death.^4^

While UV and visible light have only hundreds of microns to 1 mm of light penetration through human tissue (skin, muscle, fat), the NIR window of 650 nm to 900 nm, also known as the optical therapeutic window, is ideally suited for *in vivo* applications because of minimal light absorption by hemoglobin and water with significant penetration through human tissue reaching ~10 cm.^5^ We have previously exploited two-photon NIR activation of Feringa-type motors for inducing rapid cellular necrosis, but that technique requires large laser-generated fluxes of photons and hence the depth of penetration is shallow, ~0.5 mm, and the area of coverage is restricted to smaller-sized domains, impractical for broad clinical translation.^6^ Here we capitalize on a new type of molecular mechanical action upon cells induced by single-photon NIR light. We discovered that when we activate the vibronic mode of a cell-membrane-associated molecule, it results in the concerted whole-molecule-vibration,^7–9^ on picosecond timescales,^10^ inducing rapid cell death. Cell-associated MJH exhibiting this concerted whole-molecule VDA have a mechanical action distinct from the UV-Vis-activated unidirectional rotary molecular drills based on the Feringa and Dube designs.

In a typical molecular absorbance of photons, an individual bond or small portion of the molecule starts vibrating (Fig. 1a) or many bonds vibrate in a disconcerted manner (Fig. 1b). But there is another way to excite a molecule wherein a whole-molecule-vibration or collective vibration is achieved that is much longer-range and concerted, spreading through the entire length or width of the molecule (Fig. 1c). When vibrational and electronic modes, sometimes called the phonon and plasmon modes, respectively, are coupled, the two modes together result in vibronic coupling,^9,11^ or it can be called a molecular plasmon-phonon coupling.^8^ More specifically, by absorbance of a suitable energy of light, a molecule’s vibrational modes hybridize with the molecule’s electronic transitions to induce the vibronic mode. The vibronic mode is analogous to an ultrafast breathing mode of a molecule where the entire molecule is vibrating in unison throughout its length and/or its width because one can have a longitudinal or transverse collective vibration, respectively.^8,10^ Here we show that when a suitable molecule is cell-membrane-associated, it can rapidly compromise the integrity of the membrane in a manner and rate that no partial molecular vibration (Fig. 1a) or disconcerted vibrations (Fig. 1b) can induce. Heating a molecule through photothermal therapy can cause many vibrations in a molecule, but those vibrations are not coordinated, as shown in Fig. 1b, hence there is no concerted longitudinal or transverse vibration that is sufficient to rapidly open a cell membrane. Hence, high powers and extended times are needed in photothermal therapy to cause slower apoptotic death. Conversely, VDA of a cell-associated molecule results in rapid necrosis even at very low energies. Likewise, VDA is distinct from photodynamic therapy where the latter generates ROS, while VDA in a cell-associated molecule causes cell death that is unaffected by even large doses of ROS-inhibitors. Cyanine dyes have been used in photothermal and photodynamic therapies and they are readily accepted in biological and medicinal studies.^12–16^ However, here we use these same classical structures in what is generally 10x lower concentration and with 10-50x lower powers than often used (80 mWcm^-2^ instead of 1 to 4 Wcm^-2^), exploiting their VDA to kills cells 10-50x faster than with photothermal or photodynamic therapies. By exploiting low power NIR light for this molecular cell-killing, future use in medicinal applications holds great promise.

**Fig. 1.**
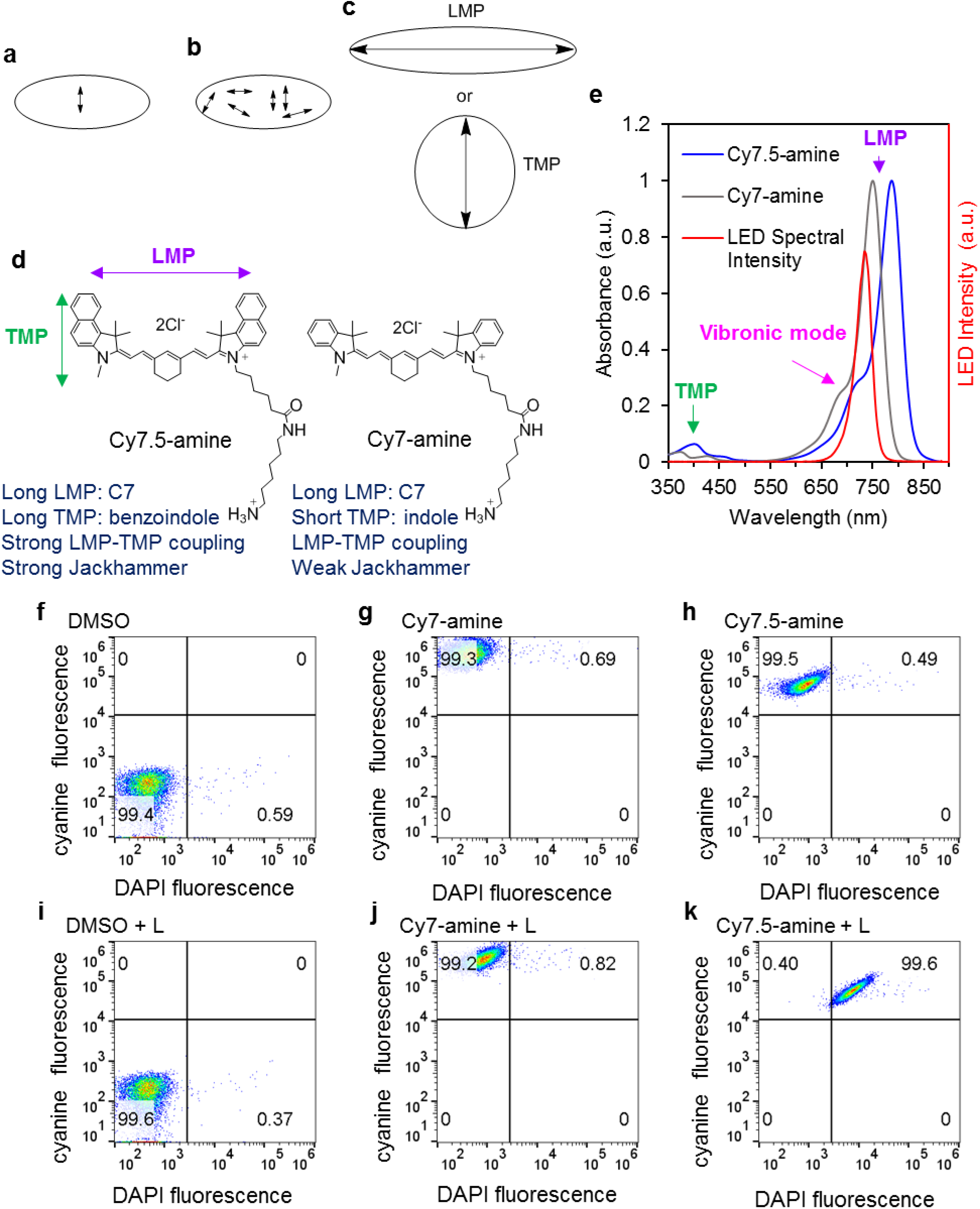
Schematics, structures, spectra, and flow cytometry. Schematic representation of bond vibrations represented by arrows using (**a**) light excitation of a single bond, (**b**) broadband light excitation of multiple bonds and (**b**) vibronic-driven action (VDA) for whole-molecule excitation with a longitudinal molecular plasmon (LMP, top) or transverse molecular plasmon (TMP, bottom). (**d)** Chemical structure of Cy7.5-amine and Cy7-amine and expected molecular plasmon modes TMP and LMP. Listed below are their structural features and translated vibrational actions. (**e**) Absorption spectra of cyanine molecules and assignment of vibrational collective oscillation (vibronic mode), LMP and TMP. Spectral intensity of the LED source and overlapped with the absorption spectrum of cyanine molecules. The spectral intensity of the LED has a λ_max_ = 734 nm and full width at half maxima (FWHM) = 35 nm. The LED light excites mostly the vibrational mode in Cy7.5-amine but not in Cy7-amine. Spectral intensity of the LED was provided by the vendor (UHP-F-730nm, Prizmatix, Israel). (**f-k**) Flow cytometry analysis. Fast A375 melanoma cell permeabilization to DAPI (DAPI enters and stains the membraned-disrupted cells but slower with viable cells) immediately upon treatment with 1 μM Cy7.5-amine or Cy7-amine excited with 730 nm NIR light (80 mWcm^-2^ for 10 min) and analyzed within ~1 min after the light treatment. (**f**) Control 0.1% DMSO. (**g**) Cy7-amine. (**h**) Cy7.5-amine. (**i**) Control 0.1% DMSO + NIR light treatment. (**j**) Cy7-amine + NIR light treatment. (**k**) Cy7.5 amine + NIR light treatment. The numbers inside the gates (four quadrants) in the flow cytometry plot represent the percentage of cells in each gate: cyanine negative and DAPI negative (left bottom), cyanine positive and DAPI negative (left top), cyanine negative and DAPI positive (right bottom), and cyanine positive and DAPI positive (top right). All the cell suspensions for this study contained 0.1% DMSO which is used to pre-solubilize the cyanine molecule at 8 mM stock solution in 100% DMSO. 10,000 cells were analyzed in each condition.

## Results

Fig. 1d-e shows the chemical structure and absorption spectra of two aminocyanines, Cy7.5-amine and Cy7-amine. Cyanine structures are characterized by an odd-numbered polyene linker connecting two nitrogen-containing heterocycles with unusual photophysical properties. The absorption spectrum of cyanines is dominated by an absorption band in the visible/NIR electromagnetic spectrum with a shoulder located at higher energy (shorter wavelength). The Cy7.5-amine, in contrast to Cy7-amine, has an additional aryl ring that increase the conjugation which cause a red-shifting of the absorption by ~40 nm relative to Cy7-amine.

Here we propose that the feature of the higher-energy shoulder next to the large absorption band in symmetrical cyanine structures results because there is coupling of a molecular plasmon (a dominant collective oscillation of electronic excitation) to a dominant collective vibrational excitation, in agreement with the suggestion in the literature that the absorption sub-bands in cyanines are primarily determined by a dominant symmetric vibration rather than a collection of single vibrations.^7^ This vibronic behavior, through the coupling of electronic and vibrational states, is a feature of the conjugated-backbone-near-symmetrical cyanines such as in Cy7-amine and Cy7.5-amine. The shoulder (λ ~ 730 nm) in the absorption spectrum of Cy7.5-amine corresponds to this collective vibrational mode (Fig. 1e). The same collective vibrational mode is present in Cy7-amine but at ~ 690 nm. The molecular plasmons in cyanines were indeed confirmed by Time-Dependent Density Functional Theory (TDDFT) calculations; these molecules can support longitudinal molecular plasmons (LMP) and transversal molecular plasmons (TMP) (Fig. 1d, 1e and Extended Data Fig. 1). The shoulder band is not the only vibronic mode present in the absorption spectrum, but instead probably the strongest in vibronic character spreading throughout the length and width of the molecule. The larger band at ~780 nm, dominantly a longitudinal charge density resonance, and the small peak at ~400 mm, with a transversal charge density resonance, and the shoulder at ~450 nm, with a short longitudinal charge density resonance, also have vibronic properties with charge density resonances typical of a molecular plasmon,^8^ as shown in Extended Data Fig. 1.

Here we selectively excite the vibronic mode in a cell-membrane-bound Cy7.5-amine using a NIR light-emitting diode (LED) at 730 nm (Fig. 1e) which results in the permeabilization of the cellular membrane to 4’,6-diamidino-2-phenylindole (DAPI). DAPI is a cell membrane impermeable dye in viable cells that mainly stains cellular DNA in membrane-disrupted cells, with induction of rapid necrotic cell death in human A375 melanoma cells (Fig. 1f-k). While 730 nm light does not excite the vibronic shoulder of Cy7-amine, it can activate the Cy7.5-amine (Fig. 1e) and permeabilize A375 cells immediately after treatment. It took ~30 s from the time the sample was irradiated to start collecting the data in the flow cytometer. For this experiment, 1 μM Cy7.5-amine, 730 nm LED at 80 mWcm^-2^ for 10 min, caused permeabilization to DAPI staining of 99.6% of the A375 cells in a cell suspension containing 2×10^5^ cells. In contrast, Cy7-amine was not able to permeabilize the cells under the same conditions. This difference between the two aminocyanines supports the notion that the 730 nm light can excite the vibronic mode (shoulder) in Cy7.5-amine (Fig. 1e) causing cellular membrane permeabilization and ultimately cell death by necrosis as seen by immediate DAPI staining. Both aminocyanines were attached efficiently to the cells as shown by flowcytometry; they were both loaded into the cell (Fig. 1g-h). It is interesting that Cy7-amine has a larger extinction coefficient (ε = 132,000 M^-1^cm^-1^) at λ = 730 nm than Cy7.5-amine (ε = 72,000 M^-1^cm^-1^) in water, yet Cy7-amine does not permeabilize the cells at comparable concentrations while Cy7.5-amine permeabilizes readily upon excitation (Fig. 1). This also suggests that a photothermal effect is not operating since Cy7-amine has a higher absorption cross-section than Cy7.5 at 730 nm. Likewise, photodynamic therapy is unlikely since Cy7 cyanines and Cy7.5 cyanines have similar yields for singlet oxygen generation.^17^ This will be further discussed below. A summary of the proposed working mechanism is described in Extended Data Fig. 2. First, the aminocyanine binding to the cells is possibly mediated by the charge on its pendant amine moiety to the negatively charged phospholipids (Extended Data Fig. 3) follow by light-activated VDA that open cell membranes. Likewise, the aminocyanine Cy5-amine was added as competitor for cell binding against Cy7.5-amine, the results showed a reduction of permeabilization activity of Cy7.5-amine (Supplementary Information Fig. S1). Extended Data Fig. 4 summarizes the optical spectra of all aminocyanines in this study and the characterization of their binding to the A375 human melanoma cells by confocal microscopy.

Consistent with the VDA proposed here, excitation of the 680 nm vibronic shoulder in Cy7-amine improves the MJH effect for opening cell membranes in A375 cells (Extended Data Fig. 5). However, the permeabilization is not as large as for 730 nm excitation of Cy7.5-amine even when exciting the 680 nm shoulder of Cy7-amine using an equal light dose of 80 mWcm^-2^ for 10 min. This might result because Cy7-amine lacks the extended aryl ring of Cy7.5-amine, limiting the TMP of the vibronic mode. Hence, Cy7-amine is a weaker MJH than Cy7.5-amine because of the lower plasmonicity of the indole in contrast to the benzoindole (Fig. 1).

Fig. 2 compares the Cy7-amine and Cy7.5-amine when exciting at 730 nm at various concentrations. It is evident that the Cy7.5-amine is much more efficient at permeabilizing cells upon the VDA with NIR light. Even at much higher concentrations of Cy7-amine (8 μM), it does not permeabilize cells as efficiently as lower concentrations of Cy7.5-amine. This is further confirmation that the excitation of the vibronic shoulder in Cy7.5-amine at 730 nm and the extension of the conjugation by the aryl rings in the benzoindoles are critical to maximize the VDA. The other example of an aminocyanine family, Cy5.5-amine and Cy5-amine, follows a similar behavior. The results suggest that there is a molecular structure/VDA intensity correlation with the molecular mechanical action (Fig. 2). The VDA increases with the length of the conjugation in the polymethine bridge and the extension of the aromatic ring at the indole.

**Fig. 2.**
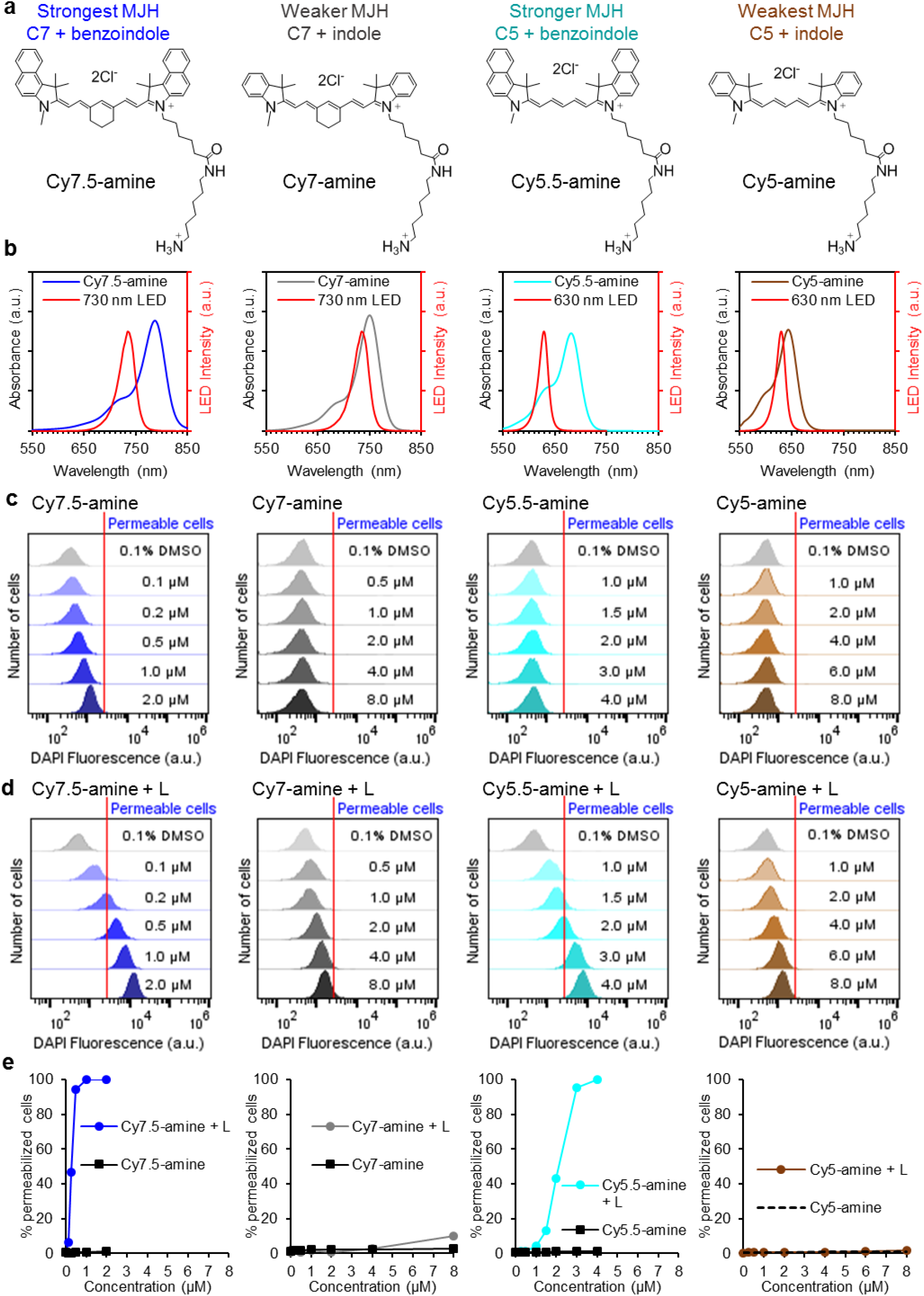
Cell membrane permeabilization dependence with the expected strength of the MJH. (**a**) Structures of MJH and classification according to their expected relative strength in the vibronic-driven action (plasmonicity) based on the extension of the indole with polycyclic aromatic hydrocarbons (PAH) and the length of the π-conjugation in the polymethine bridge. (**b**) The absorption spectra of each MJH overlaid with the specific LED light that was used for illumination in this experiment. (**c**) Flow cytometry analysis to measure the permeabilization of A375 cells in the presence of each MJH without illumination. The red line represents the gating to discriminate between DAPI negative and positive cells (permeable). (**d**) Flow cytometry analysis to measure the permeabilization of A375 cells in the presence of each MJH with specific LED illumination for each cyanine. The red line represents the gating to discriminate between DAPI negative and positive cells (permeable). (**e**) Plots of the percentage of permeabilized cells, the numbers are extracted from the flow cytometry analysis. 10,000 cells were analyzed in each concentration. The incubation of the MJHs with the cells was 30 min. The light-treated samples were illuminated with the same light dose of 80 mWcm^-2^ for 10 min. The shifting on the DAPI fluorescence intensity for the Cy7.5-amine before illumination correspond to the fluorescence emission from Cy7.5-amine which is excited with the same laser that excites DAPI (λ_ex_ = 405 nm). Extended Data Fig. 3 supports the conclusion that this observation correlates with the confocal microscopy data. This fluorescence level from Cy7.5-amine can be regarded as background fluorescence and not cell membrane permeabilization. This main factor was considered to draw the position of the gating (red line) to discriminate between DAPI positive cells and DAPI negative cells.

Fig. 3 shows the confocal microscopic permeabilization of the cells over time using 630 nm light treatment of Cy5.5-amine and Cy5-amine where the vibronic band in Cy5.5-amine is accessed while only weakly accessed in Cy5-amine (Fig. 2b). The cell permeabilization was done in the presence of the cell-membrane-targeting DiD dye under the confocal microscope. At 4 min of laser excitation, the cells are already permeabilized; the DAPI intensity (cell permeabilization) is 2x relative to the initial and 13x higher at 10 min. The result show that Cy5.5-amine is a stronger MJH than Cy5-amine (2.5x DAPI intensity increase at 10 min) which is consistent with the flow cytometry analysis in Fig. 2c-e. The DiD dye alone showed a weak effect on cell permeabilization (1.5x DAPI intensity increase at 10 min), as expected because it is analogous to Cy5, a MJH with a weak VDA. In all non-illuminated controls, the DAPI intensity remains at 1x relative to the initial conditions.

**Fig. 3.**
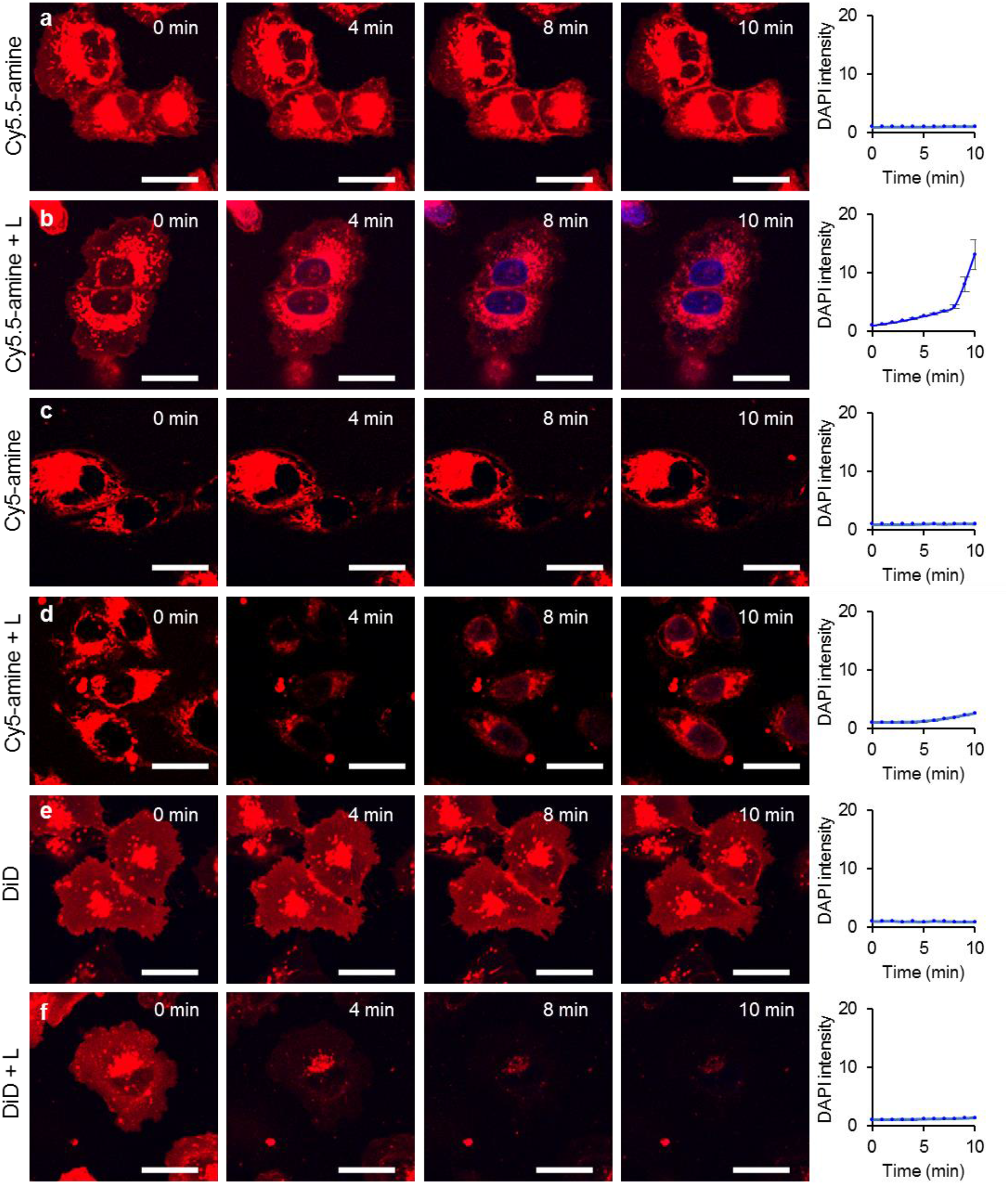
Cell membrane permeabilization of A375 cells over time while the cells were irradiated under the confocal microscope. The permeabilization of DAPI into the cells was recorded as a function of time (rightmost column). (**a**) Cells in the presence of 4 μM Cy5.5-amine without laser irradiation. (**b**) Cells in the presence of 4 μM Cy5.5-amine with 640 nm laser irradiation. (**c**) Cells in the presence of 4 μM Cy5-amine without laser irradiation. (**d**) Cells in the presence of 4 μM Cy5-amine with 640 nm laser irradiation. (**e**) Cells in the presence of cell-membrane-targeting 4 μM DiD dye without laser irradiation. (**f**) Cells in the presence of 4 μM DiD dye with 640 nm laser irradiation. The irradiation times are shown in each image. For the activation of the MJH effect, the cells were irradiated at λ_ex_ = 640 nm, 25% power. The pictures were recorded every 1 min at low photoactivation powers and short exposures (λ_ex_ = 640 nm, 5% power, 2.1 s per a single frame) to maintain as intact as possible the non-irradiated controls. The incubation of the cyanines with the cells before irradiation was 30 min. Representative confocal images of each condition are shown. The quantification on the rightmost is the average of 14 cells (n = 14), the error bars are the standard errors. All scale bars = 25 μm.

To rule out a photothermal effect, the temperature of the media was measured during the NIR light exposure when the treatment was done at room temperature versus when done in an ice bath (Extended Data Fig. 6) surrounding the cell culture dish. The temperature remained constant at 20 °C and 2 °C when the treatment was done at room temperature and in an ice bath, respectively. In Extended Data Fig. 6g, the cell permeabilization is not affected by the lower temperature. Thus, the photothermal effect is not responsible for the necrosis seen in these cells.

To confirm that a photodynamic ROS generation is not responsible for the necrosis, the permeabilization of A375 melanoma cells was repeated in the presence of ROS scavengers (Extended Data Fig. 7a-c). Neither *N*-acetyl-cysteine (NAC, 10 mM), thiourea (TU, 100 mM) or sodium azide (SA, 2.5 mM) were able to retard the permeabilization of the cells. Further, the ROS is not responsible for the permeabilization of the cells; in Extended Data Fig. 7d the permeabilization conditions were tuned to lower illumination time where the scavengers might quench ROS more effectively. The results show that none of the ROS scavengers used (NAC, TU, SA, methionine or vitamin C) slowed the permeabilization of the A375 cells. This suggests that ROS is not responsible for the permeabilization of the cells. Extended Data Fig. 8 shows that singlet oxygen production is also not responsible for the permeabilization of the cells.

To confirm that the permeabilized cells to DAPI are indeed dead, we cultured the cells after the treatment (with 2 μM Cy7.5-amine and illumination for 10 min with 730 nm light at 80 mWcm^-2^) and quantified cell death by crystal violet and clonogenic assays. Extended Data Fig. 9 shows that the permeabilized cells by light-activated Cy7.5-amine were dead nearly at the same quantity, 99.9%, as was observed in flow cytometry (Fig. 1). The clonogenic assay shows that 100% of the cells were eradicated when using 0.5 μM Cy7.5-amine and illumination for 10 min with 730 nm light at 80 mWcm^-2^.

The FDA-approved indocynine green (ICG) bearing alkylsulfonate addends was also tried; it has a vibronic absorption shoulder at 730 nm, as shown in Supplementary Information Fig. S2. The sulfonates will not interact with phospholipids in lipid bilayers. ICG did not permeabilize the cells under the same conditions in which Cy7.5-amine was tested. Taken together, this suggests that the effective cell killing of Cy7.5-amine is a synergy of the amine pendant for rapid cell association followed by VDA. This presumably causes an energy transfer from these whole-molecule excited structures to the cellular membrane, ultimately disrupting the lipid bilayer, permitting cell membrane permeabilization.

The MJH Cy7.5-amine was applied to treat murine (B16-F10) and human (A375) melanoma tumors in mice (Fig. 4). The temperature of B16-F10 tumors in C57BL/6 mice while under Cy7.5-amine with light treatment (150 mWcm^-1^ for 5 min) increased ~5 °C and this was not different than the control with 0.1% DMSO and light (Fig. 4b-c). The size of the B16-F10 tumors was significantly reduced using a dose of 8 μg of Cy7.5-amine in 50 μL PBS solution intratumorally and illumination with 730 nm LED at 150 mWcm^-1^ for 5 min (Fig. 4 d) In the treatment of A375 tumors in nude mice, the conditions were optimized, and we found that 300 mWcm^-2^ of 730 nm LED for 5 min in combination with a intratumoral dose of 8 μg of Cy7.5-amine was sufficient to achieve a survival rate of 60% at day 120 of the study and 50% of the mice became tumor free.

**Fig. 4.**
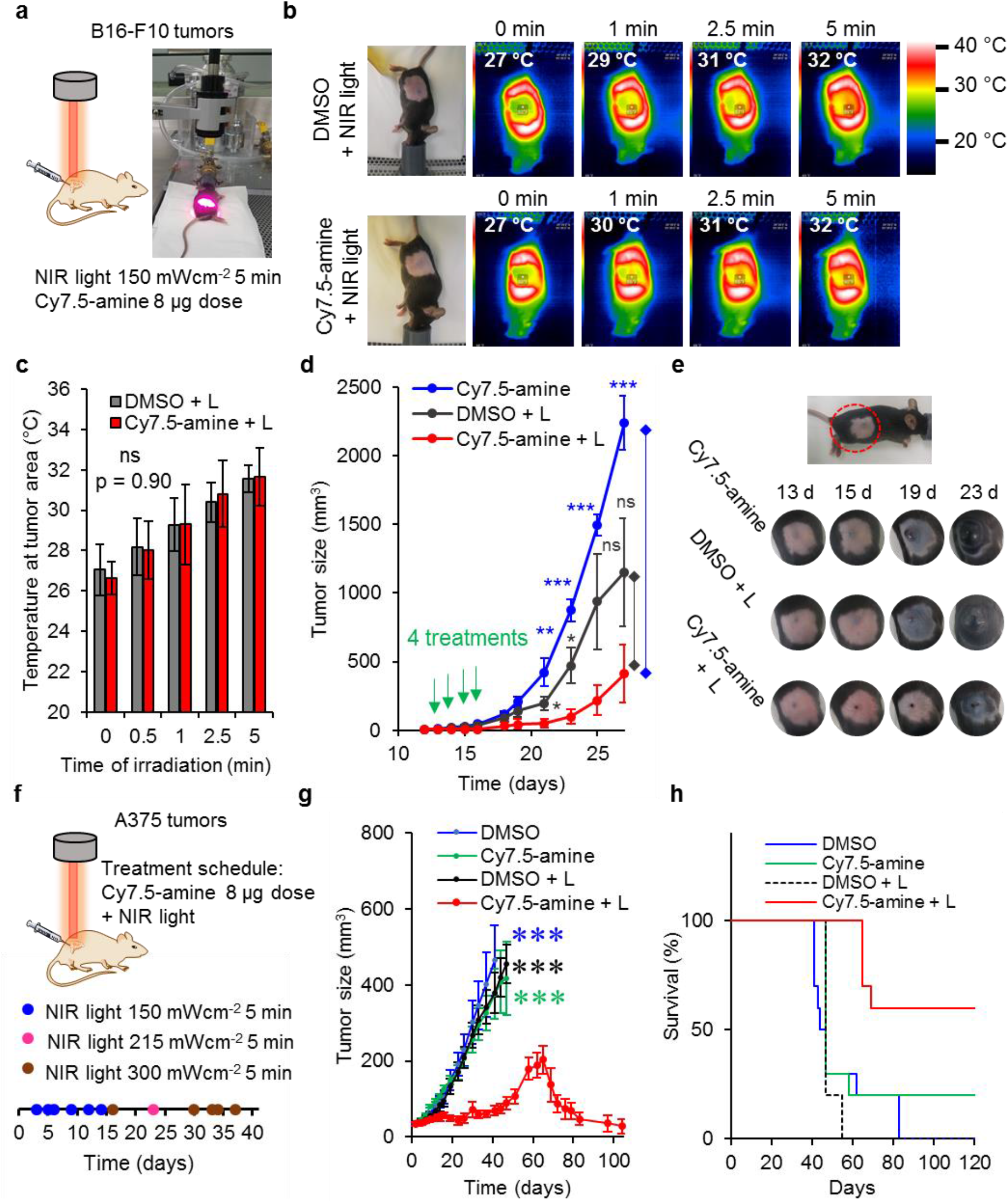
Therapeutic effect of MJH Cy7.5-amine in the treatment of tumors in mice. (**a**) Pictures of the set up and the conditions to treat B16-F10 melanoma tumors in C57BL/6 mice. The Cy7.5-amine was applied by intratumoral injection of 50 μL solution containing 0.16 mg mL^-1^ in PBS solution with 0.1 % DMSO. 0.1 % DMSO in PBS is use as a control. (**b)** Recording of the temperature on the mouse skin using a NIR camera while being treated with Cy7.5-amine or DMSO and irradiated with a 730 nm LED at 150 mWcm^-2^ for 5 min. The inset in each picture shows the temperature on the tumor area. On the top of the picture is the time of illumination. (**c**) Temperature of the tumor area as a function of time. There is little difference with respect to the control. Four independent measurements n = 4. Error bars are the standard deviations. (**d**) Effect of the treatment on the size of the B16-F10 tumors as a function of time. Number of mice per group, n = 4. Error bars are the standard errors. (**e**) Representative figure of mice with B16-F10 tumors and the response to treatment. The treated tumors become dry with a necrotic spot upon treatment with Cy7.5-amine + 150 mWcm^-2^ for 5 min of 730 nm LED. (**f**) Schematics of the conditions and treatment schedule on A375 human melanoma tumors in nude mice. The Cy7.5-amine was applied by intratumoral injection of 50 μL solution containing 0.16 mg mL^-1^ in PBS solution with 0.1 % DMSO. 0.1 % DMSO in PBS is use as a control. Three doses of light were applied in combination with the solutions (Cy7.5-amine or DMSO). (**g**) Effect of the treatment on the size of the A375 tumors as a function of time. Number of mice per group, n =10. We learned in the process that 150-215 mWcm^-2^ produced a mild therapeutic effect in A375 tumor models. With 300 mWcm^-2^ a strong effect on the regression of the tumor size was observed. The drop at 60 days in the Cy7.5-amine + L group was due to the exclusion of the euthanized mice. Error bars are the standard errors. *t*-test, two-tail, * p < 0.05, ** p < 0.01, *** p < 0.001. Statistical significance p < 0.05. (**h**) Survival of mice with A375 tumors under the various treatments (n = 10 mice per group). 60% of the (6 out of 10) mice survived and 50% are tumor free in the Cy7.5-amine + light (L) group. Two mice (20%) in the Cy7.5-amine group survived.

## Conclusion

VDA in low concentrations of MJH is an effective method to permeabilize cancer cell membranes and cause high efficiency necrotic cell death using low intensities of NIR light (12 J/cm^2^, 80 mW/cm^2^ for 2.5 min). With an IC50 = 250 nM, VDA of Cy7.5-amine is well-suited for rapid clinical translational study in cancer treatment. We have identified an easily NIR-accessed molecular plasmon (vibronic coupling) in a small organic molecule that is biocompatible, stable in water, and highly active for the permeabilization of cell membranes. There is no need to electrochemically add or remove electrons to these small molecules to make them vibronic-mode active. This demonstrates a new mechanical action at the molecular scale to eradicate cells or also possibly treat cells using adjuvants. We are now identifying and synthesizing other small molecules that can accentuate this combination of features for cell binding and VDA. Other applications could include the selective regulation of active sites in enzymes, modulation of protein channels, or regulation of structure/function of supramolecular biological assemblies with pharmacological implications.

## Methods

### Cyanine molecules

Cy7.5-amine, Cy7-amine, Cy5.5-amine and Cy5-amine were purchased from Lumiprobe Corp. (Maryland, USA). DiD and DiR cyanines were purchased from Biotium (Fremont, CA).

### LED illumination systems

The 730 nm LED (model UHP-F-730) and 630 nm LED (model UHP-F-630) were purchased from Prizmatix, Israel. The 680 nm LED and 740 nm LED were custom made and purchased from Keber Applied Research Inc. (Ontario, Canada).

### Culture of human melanoma A375 cells

A375 cells were obtained from ATCC (CRL-1619). Cells were cultured in 10 cm polystyrene tissue culture treated dish (Corning) containing DMEM with L-glutamine, 4.5 g/L glucose, and sodium pyruvate (Corning Inc. 10013CV) and supplemented with 10% FBS (Corning, 35010CV), 1X MEM vitamin solution (Gibco, 11120052), 1X MEM non-essential amino acid solution (Gibco, 11140050) and penicillin/streptomycin. Typically, 0.5-1 million cells are inoculated per dish, and culture for 3-4 days in incubator at 37 °C and 5 % CO_2_, then transferred to a new dish when confluency reached nearly 95-100%. For the passage step, cells are detached with 0.05 % trypsin-EDTA (Gibco, 25-300-054).

### Culture of mouse melanoma B16-F10 cells

Mouse melanoma B16-F10 cells were obtained from the ATCC (CRL-6475) and cultured in 10 cm polystyrene tissue culture treated dish (Corning) containing DMEM with 4.5 g/L glucose (Gibco, 11960-044) and supplemented with 10% FBS (SAFC Industries-Sigma-Aldrich, 12303C), 2X (10 mL) MEM vitamin solution (Corning, 25-020-Cl), 1X (5 mL) non-essential amino acid (NEAA) mixture (Lonza, 13-114E), 1X (5 mL) of L-glutamine (Lonza, 17-605E), and 1X (5 mL) of penicillin/streptomycin (Hyclone, SV30010). Typically, 0.5-1 million cells were inoculated per dish, and cultured for 2-3 days in incubator at 37 °C and 5 % CO_2_, then transferred to a new dish when confluency reached nearly 95-100%. For the passage to a new culture dish, cells are detached with 0.05 % trypsin-EDTA (Gibco, 25-300-054).

### Cell permeabilization and flow cytometry analysis

A375 cells were cultured as described before. One day before the treatment, cells were inoculated at 5 million cells per dish (10 cm polystyrene tissue culture dish). The cells were harvested using 0.05% trypsin-EDTA (Gibco, 25-300-054), then the cells were counted and were adjusted to a cell density of 2×10^5^ cells/mL in DMEM media with L-glutamine, 4.5 g/L glucose, and sodium pyruvate (Corning Inc. 10013CV) and supplemented with 10% FBS (Corning, 35010CV), 1X MEM vitamin solution (Gibco, 11120052), 1X MEM non-essential amino acid solution (Gibco, 11140050) and penicillin/streptomycin. 1 mL of this cell suspension containing 2×10^5^ cells was used in each treatment. In a 1.5 mL Eppendorf tube, 1 μL of stock solution containing 2 mM Cy7.5-amine (or other cyanine molecule or other concentration) in DMSO (Fisher, 99.7%) was placed in the bottom of the tube, then 1 mL of the cell suspension was added into the tube to get final concentration of 2 μM of Cy7.5-amine containing 0.1% DMSO and 2×10^5^ cells. The mixture was then incubated at 37 °C and 5% CO_2_ for 30 min. Then, 1 μM DAPI was added into the cell suspension. Then, the cells suspension was transferred to a 35 mL polystyrene tissue culture dish and immediately the cells were treated under the light beam of NIR light of 730 nm at 80 mW/cm^2^ (or adjusted powers down to 20 mW/cm^2^) for 10 min (or adjusted illumination times down to 30 s) using LED light source (Prizmatix, UHP-F-730, Israel) which covers the entire dish. The spectral intensity of the LED is shown in Fig. 1e. While the cells were treated, the dish was placed on top of an aluminum block painted black, so that the excess NIR light and that was not reflected back into the cell suspension while the aluminum block acts as a heat-sink, maintaining a constant temperature in the dish during the irradiation. The instrument for flow cytometry analysis (SONY, MA900 Multi-Application Cell Sorter using the LE-MA900 FP-Cell Sorter Software) was already set up and calibrated by the time the light treatment was finished. Therefore, as soon as the 10-min light treatment was completed, the cell suspension was rapidly transferred from the 35 mm dish to a flow cytometry tube and the cells were analyzed for DAPI permeabilization and Cy7.5-amine binding. It took ~30 s to load the sample and to start the analysis. Therefore, the permeabilization of cells was measured as DAPI positive cells and occured immediately due to the membrane permeabilization caused by Cy7.5-amine excitation with the 730 nm NIR light. The flow cytometry data was analyzed using FlowJo software. The light intensity was measured using an Optical Power Meter from Thorlabs, sensor model S302C and console model PM100D.

### Confocal microscopy

For confocal microscopy, 80,000 A374 cells were inoculated in a glass bottom dish (IBIDI, μ-dish 35 mm high glass bottom) in DMEM media with all supplements as described before and culture at 37 °C and 5% CO_2_ for 1 day. On the day of the confocal microscopy analysis, the DMEM media was removed and the cells were washed twice with PBS buffer. Then a mixture of fresh DMEM media without phenol red and with the MJH was added to the cells. The DMEM without phenol red was from Gen Clone, product number 25-501C, and it was supplemented with 10% FBS (Corning, 35010CV), 1X MEM non-essential amino acid solution (Gibco, 11140050) and penicillin/streptomycin. The cells were incubated in the presence of the MJH at 37 °C and 5% CO_2_ for 30 min. Then the cells were imaged and photoactivated in the confocal microscope model Nikon A1-Rsi (Rice University Shared Equipment Authority).

### Temperature measurements

The permeabilization of the cells and flow cytometry analysis was conducted as described above. The temperature of the cell suspension was measured using the temperature probe (Model SC-TT-K-30-36-PP; Omega Engineering, Inc.) immersed in the media. The same was repeated having the cell suspension on top of an ice bath, and the temperature of the cell suspension recorded in the same way during the NIR light illumination. The temperature of the media stays constant at room temperature of ~20 °C upon illumination of the media with the 730 nm LED light at 80 mW/cm^2^ for 10 min. There is only a minor temperature increase of 0.4 °C that is attributed to the light illumination absorption by the media components. Similarly, on ice the temperature of the media only increased 0.6 °C due to the illumination by the 730 nm LED on the media.

### ROS scavenger experiments

The permeabilization of the cells and flow cytometry analysis was conducted as described before. But in this case, ROS scavengers were added into the cells suspension and incubated for 1.5-2 h at 37 °C and 5 % CO_2_ before any treatment to allow the antioxidants to interact first and protect the cells.

### Crystal violet cell viability assay

The crystal violet assay was used to measure the cell viability. The principle of this method is that the viable cells adhere to the surface of the cell culture dish and keep growing and remain attached through the standard cell culture conditions during a period of 1 to 2 days and through the staining conditions in the assay. In contrast the dead cells do not adhere to the surface of the cell culture dish, do not grow, and detach easily during the manipulation steps during the assay, which includes removal of media and exchange with fresh media and washing steps with PBS buffer. For specific details, A375 cells were harvested and counted, and then 20,000 A375 cells per well were added in 24 cell culture well plate (Corning) and cultured for 1 day at standard incubation conditions of 37 °C and 5% CO_2_. The cells were treated in four experimental groups (4 samples per group): group 1) 0.1% DMSO, group 2) 0.1% DMSO + NIR light treatment, group 3) 2 μM Cy7.5-amine, and group 4) 2 μM Cy7.5-amine + NIR light. The treatments with 0.1% DMSO or 2 μM Cy7.5-amine were done by adding those respective concentrations to the cells in the media and then incubation for 60 min. Immediately after the incubation, the cells in the groups with “+ NIR light”, were treated with 730 nm light at 80 mW/cm^2^ for 10 min. After the treatment with NIR light, the media in all the groups was removed and fresh media was added. Then, the cells were incubated for 2 days at 37 °C and 5% CO_2_. At the end of the incubation, the media was removed and the cells washed with 500 μL of PBS once. Then the cells were stained with 500 μL of 0.5% w/v crystal violet solution in methanol/water (1:1) for 5 min. Then the crystal violet was removed, and the excess of crystal violet was washed with water. The cells contained in the 24 well plate were dried at room temperature. Then, the crystal violet in each well was solubilized in 500 μL of 3.3% v/v acetic acid in water and the total crystal violet recover in this acidic solution. Lastly the crystal violet was quantified by its absorbance at 570 nm. The cell viability was calculated from the absorbance relative to the absorbance in the cells without any treatment, cell without treatment were normalized to 100% cell viability.

### Clonogenic assay

A375 cells were seeded in 35 mm cell culture dishes at predetermined densities to allow for an approximately equal number of resultant colonies. The next day, cells were treated with Cy7.5-amine at variable concentration and with or without 730 nm light at 80 mWcm^-2^ for 10 min. The cells were incubated with Cy7.5-amine for 50 min before the illumination, the media was replaced with fresh media after the illumination and cells were cultured for 6 days to allow for colony formation. Cells were then washed once with PBS and fixed-stained in a 0.5% (w/v) crystal violet in methanol/water solution (1:1) during 10 min. The excess of crystal violet was washed off with water, the plates were dried at room temperature and then the colonies were counted using ImageJ software version 1.52a, and the survival fraction was determined. All treatments were in triplicate.

### ROS measurements using H2DCF-DA

H2DCF-DA (2’,7’-dichlorodihydrofluorescein diacetate) is a cell permeant reagent. It is deacetylated by cellular esterases to form 2’,7’-dichlorodihydrofluorescein (H2DCF), a non-fluorescent compound, which is rapidly oxidized in the presence of ROS into 2’,7’-dichlorofluorescein (DCF). DCF is highly fluorescent and is detected with excitation / emission at 488 nm / 535 nm. A375 cells in suspension containing 2×10^5^ cells mL^-1^ were first prepared in DMEM media without phenol red. Then the cells were incubated for 30 min at 37 °C with Cy7.5-amine (or the other cyanines) typically at 2 μM concentration in the media. Then, H2DCF-DA (Sigma-Aldrich) was added to cells suspension in media to the final concentration of 5 μM (the stock of H2DCF-DA was at 5 mM in DMSO stored at −20 °C). Then transfer the cells to a 96 well plate, 100 μL to each well. Typically, it took 15 min to transfer all the samples and all the controls into the 96 well plate, 6 repetitions per each treatment condition: cyanine + light, cyanine only, DMSO + light, DMSO only, cells only, media only. Then, immediately after the cells were treated with NIR 730 nm light at 80 mWcm^-2^ for 10 min. From the time the H2DCF-DA was added into the cell suspension to the time the cells were treated there was in total a 20 min incubation. After the light treatment, the DCF fluorescence intensity was measured immediately after the light treatment using a 96 well plate reader at λ_ex_ = 488/9 nm and λ_em_ = 535/20 nm. The measurements were normalized with respect to the fluorescence intensity in the media only.

### Singlet oxygen measurements

For the measurement of singlet oxygen, the molecular probe DPBF (1,3-diphenylisobenzofuran) was used which decomposes in the presence of singlet oxygen and this was detected by the change in the absorbance of DPBF at 410 nm. DPBF was freshly prepared for every experiment by dissolving DPBF in methanol at 1 mM stock solution. Then dilute the DPBF in methanol and adjust the dilution volume to get an absorbance of ~1.0 at 410 nm. For this purpose, typically 170 μL of the 1 mM DPBF stock in methanol was diluted in 2830 μL of methanol. Then to this mixture was added 1 μL of cyanine stock solution (8 mM cyanine solution in DMSO stored at −20 °C) to get a final concentration of 2.6 μM of cyanine. Then immediately after preparing the mixture, the solution was transferred to a clean spectrophotometer quartz cuvette, and the solution was irradiated with 730 nm LED light at a power intensity of 80 mWcm^-2^. The power intensity was calibrated to the distance at the top of the liquid on the quartz cuvette. The sample was irradiated every 30 s, and then absorption spectrum was recorded in between every irradiation interval until a total of 10 min of irradiation was accumulated.

### *In vivo* studies

All animal studies were approved by the Institutional Animal Care and Use Committee (IACUC) of the University of Texas MD Anderson Cancer Center (Houston, TX). In our studies we used 7-8 weeks old female C57BL/6J mice (Jackson Laboratories, strain #000664) or 7-8 weeks old athymic female nude (nu/nu) mice from Envigo/Harlan labs.

### Injection of B16-F10 cells subcutaneously in C57BL/6J mice to generate the melanoma tumors

The B16-F10 cells were cultured as described before. Cells were harvested from sub-confluent plates, ~90%, and fresh media was added to the cells the day before harvesting. The cells were harvested using 0.05 % trypsin-EDTA (Gibco, 25-300-054). The harvested cells were re-dispersed in DMEM media without supplements at 1×10^6^ cells mL^-1^. The cell suspension was kept in ice. Then 100 μL of cells were injected per mouse (this was 100,000 cells per mouse) subcutaneously in the right flank of a 7-8 week old female mouse (C57BL/6J), in which the hair in the right flank was previously depilated using a shaver. The tumors were allowed to grow for 12 days counting from the day of cell injection. And at day 12 the hair of the mouse was removed using hair remover cream (Nair Hair Remover Lotion). For this purpose, a drop of the cream was placed on the skin, on top of the area where the tumor was injected. The mice were anesthetized using isoflurane while the hair remover cream was applied. Starting at day 12 the tumors were measured using a caliper. The tumors can be observed as a black spot (due to the melanin present in the B16-F10 cells) under the skin after the cream depilation. The typical volume of the tumors at ~15 days was ~25 mm^3^. The volume of the tumor was calculated as: (1/2) x length x width x height. When the height was not possible to measure in the case of the tumors which were too small (usually < 100 mm^3^), then the tumor volume was calculated as: (1/2) x length x width^2^.

### Preparation of fresh solution of 200 μM Cy7.5-amine and 2.5% DMSO for *in vivo* studies

In the day of treatment, fresh solution of 200 μM Cy7.5-amine was prepared by diluting in PBS buffer the 8 mM Cy7.5-amine stock in DMSO, the final dilution contained 2.5% DMSO. As control 2.5% DMSO in PBS buffer was used.

### Treatment of B16-F10 tumors with Cy7.5-amine and NIR 730 nm light

The tumors were treated at day 13 counting from the day of cell injection. Mice were divided in 3 groups: 1) Cy7.5-amine only (mice per group, n = 4), 2) 2.5% DMSO + Light (n = 5) and 3) Cy7.5-amine + light (n =5). The day of treatment, fresh solutions (200 μM of Cy7.5-amine in PBS and controls 2.5% DMSO in PBS) were prepared as described before. The mice were anesthetized with isoflurane using a vaporizer. Then, each mouse was injected with 50 μL of 200 μM Cy7.5-amine solution in PBS or 2.5% DMSO, intratumorally. Then mice were kept for 30 min in the cages to let the Cy7.5-amine solution or DMSO solution interact with the tumors. Then, after the 30 min of incubation, the mice were treated (under anesthesia, using isoflurane) with 730 nm LED light source from Prizmatix applying a power intensity of 150 mWcm^-2^ for 5 min. The light intensity was measured using an Optical Power Meter from Thorlabs, sensor model S302C and console model PM100D. While under light treatment the temperature at the tumor area was measured using an IR thermal camera (Model: Compact Seek Thermal for Android. Seek Thermal, Inc. Santa Barbara, CA). When the treatment was finished the mice were put back into the cages and housed in the animal facility. The treatment was repeated once daily for 4 days. The tumor sizes were measured every day starting the day of hair removal with cream. The tumors were measured using a caliper. Then after the 4 treatments, the tumors were measured every other day.

### Injection of A375 cells subcutaneously in athymic nude mice to generate the human melanoma tumor model

The A375 cells were culture as described before. Cells were harvested from sub-confluent plates,~90%, and fresh media was added to the cells the day before harvesting. The cells were harvested using 0.05 % trypsin-EDTA (Gibco, 25-300-054). The harvested cells were re-dispersed in DMEM media without supplements at 50×10^6^ cells mL^-1^. The cell suspension was kept in ice. Then 100 μL of cells were injected per mouse (this was 5 million of cells per mouse) subcutaneously in the right flank of 7-8 weeks old female athymic nude mouse. Starting at day 2 the tumors were measured using a caliper. The typical volume of the tumors at day 2 was ~33 mm^3^. The volume of the tumor was calculated as: (1/2) x length x width x height. When the height was not possible to measure in the case of the tumors that were too small (usually < 100 mm^3^), then the tumor volume was calculated as: (1/2) x length x width^2^.

### Treatment of A375 tumors with Cy7.5-amine and NIR 730 nm light

The tumors were treated at day 3 (since 5 million initial cells were injected per tumor site) when the average tumors size was 35 mm^3^. Forty mice were divided in 4 groups (number of mice per group, n = 10): 1) Cy7.5-amine only 2) 2.5% DMSO only, 3) 2.5% DMSO + Light and 3) Cy7.5-amine + light. The day of treatment, fresh solutions (200 μM of Cy7.5-amine in PBS and controls 2.5% DMSO in PBS) were prepared as described before. The mice were anesthetized with isoflurane using a vaporizer. Then, each mouse was injected with 50 μL of 200 μM Cy7.5-amine solution in PBS or 2.5% DMSO, intratumorally. Then mice were kept for 25 min in the cages to let the Cy7.5-amine solution or DMSO solution interact with the tumors. Then, after the 25 min of incubation, the mice were treated (under anesthesia, using isoflurane) with 730 nm LED light source from Prizmatix applying a power intensity of 150 mWcm^-2^ for 5 min (other power intensities were 210 mWcm^-2^ for 5 min and 300 mWcm^-2^ for 5 min as described in the treatment schedule in Fig. 3f). The light intensity was measured using an Optical Power Meter from Thorlabs, sensor model S302C and console model PM100D. When the treatment was finished the mice were put back into the cages and housed in the animal facility. The tumors sizes were measured every other day using a caliper in the first 20 days of the study and then in the late stage of the study twice a week.

### Time-Dependent Density Functional Theory Analyses (TDDFT)

Starting from a ground-state DFT calculation to obtain the energies of all the occupied levels, the absorption spectra were calculated by TDDFT using the Liouville-Lanczos approach as coded in the program Quantum Espresso.^18,19^ The charge density responses were visualized using the software VESTA.^20^

## Acknowledgements

J. M. T. acknowledges financial support from Nanorobotics, Ltd., the Discovery Institute, and the Welch Foundation (C-2017-20190330). We thank Dr. Ana Santos for helpful discussions. We thank Prof. Han Xiao at Rice University for kindly hosting C.A.O. and sharing his laboratory to culture the A375 cancer cells. We thank Carter Kittrel for helpful discussions. J.M.S. acknowledges the Lannater and Herb Fox Professorship.

## Author contribution

The idea to use the VDA to permeabilize cell membranes was suggested by C.A.O and discussed with J.M.T. C.A.O conducted all the experiments to demonstrated the permeabilization of cells by molecular vibrations, flow cytometry studies, confocal microscopy, *in vivo* studies, the temperature measurements, ROS measurements, and crystal violet assay, all these under supervision of J.M.T. C.A.O designed and conducted the *in vivo* experiments. A. C. injected the cancer cells to generate the tumors and assisted monitoring the mice. R.R and J.F.M. oversaw the *in vivo* experiments, provided the mice, resources and stablished B16-F10 tumor model. D.G.A. and J.M.S. conducted the TDDFT calculations of molecular plasmons in Cy7.5-amine. C.A.O. and J.M.T. wrote the manuscript. All authors read and approved the manuscript.

## Author information

Reprints and permissions information is available at www.nature.com/reprints. C.A.O is currently a research scientist at Rice University and was a former subcontractor at Nanorobotics Ltd. Readers are welcome to comment on the online version of the paper. Correspondence and requests for materials should be addressed to

## Conflict of interest

Rice University owns intellectual property on the use of MJH coupled with VDA for permeabilization of cell membranes. C.A.O. is a former subcontractor to Nanorobotics Ltd., the possible licensee of this technology from Rice University. J.M.T. is a stockholder in Nanorobotics Ltd., but not an officer, director or employee. Conflicts are mitigated through regular disclosure to and compliance with the Rice University Office of Sponsored Programs and Research Compliance. The authors declare no other conflicts.

## Data Availability

The datasets generated during and/or analyzed during the current study are available from the corresponding author on reasonable request.

---

**Extended Data Fig. 1.**
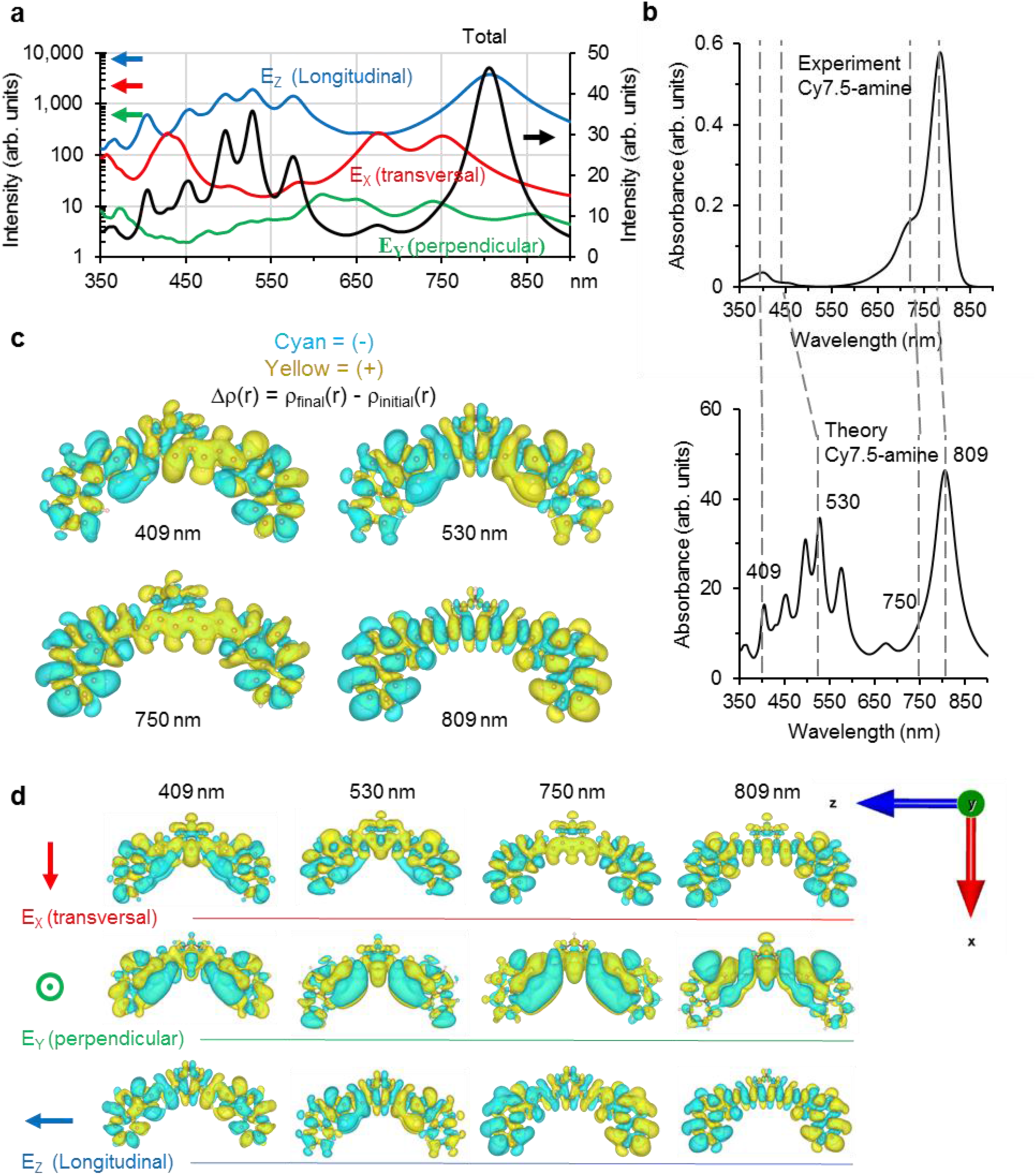
Calculated TD-DFT absorption spectrum and induced charge density plots of the molecular plasmons in Cy7.5-amine. (**a**) Total and partial, by the orientation of the electric field component (Ei), absorption spectra calculated by time-dependent density-functional theory (TDDFT) using the Lanczos approach. The electric field is used to simulate the optical excitation of the Cy7.5-amine. The partial components of the spectrum are oriented along the transversal molecular plasmon resonance (red), longitudinal (blue) and perpendicular (green) axis of Cy7.5-amine. (**b**) Absorption spectra comparison between the experimental (top) and the TDDFT calculation (bottom). The dashed lines represent the position of the wavelengths at which the induced charge density maps were calculated for molecular plasmon resonances. The experimental shoulder at 730 nm for the vibronic mode in Cy7.5-amine is observed at 750 nm in the theoretical transversal component of the spectrum, but it is less obvious in the total spectrum. (**c**) Total induced charge densities [Δρ(r)] at 409, 530, 750 and 809 nm wavelengths for molecular plasmon resonance. (**d**) Induced charge densities [Δρ(r)] by electric field (Ei) components at 409, 530, 750 and 809 nm wavelengths oriented along the transversal, longitudinal and perpendicular axis of Cy7.5-amine. The vectors on the rightmost represent the orientation of the electric filed components. The long alkyl-amine arm in Cy7.5-amine structure was not included in the electronic structure calculation because it has negligible contributions to the conjugation of the core structure. Instead, a methyl group was substituted for the long alkyl-amine in Cy7.5-amine.

**Extended Data Fig. 2.**
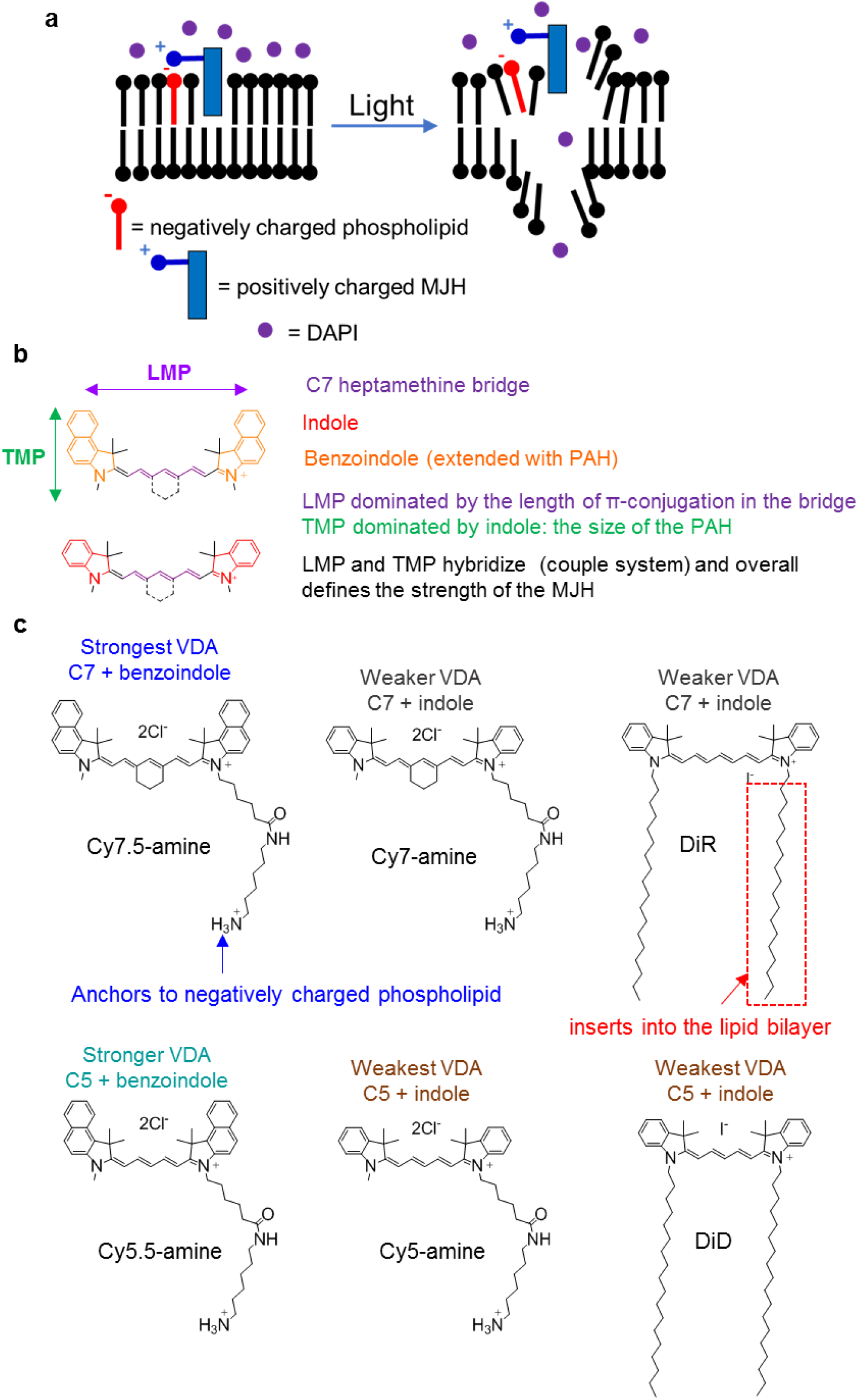
Molecular jackhammer (MJH) model and summary of structures used in this study. (**a**) General proposed working mechanism of MJH opening cell membranes. (**b**) Important MJH structural elements. LMP = longitudinal molecular plasmon. TMP = transversal molecular plasmon. The strength of the molecular plasmon (VDA) is expected to be proportional to the length of the π-conjugation. The π-conjugation can be increased in two ways: 1) increasing the length of polymethine bridge and 2) increasing the size of the polycyclic aromatic hydrocarbon (PAH) fused to the indole. The purple color is to highlight the polymethine bridge. The cyanines are named by the number of carbons in the polymethine bridge, in the example it is C7. The red color is to highlight the structure of the indole, and the orange color is for the benzoindole. The heptamethine bridge (C7) can be chemically conjugated with indole to form Cy7 or with benzoindole to form Cy7.5. These structural elements, polymethine and indole or benzoindole, hybridize to from a coupled system with the molecular plasmon-dominated longitudinally by the polymethine bridge (LMP in purple) and transversally by the indole or benzoindole (TMP in green). However, these structures are hybridized and the electronic conjugation of the benzoindole influences the polymethine bridge and *vice versa.* (**c**) Summary of structures in this study. The observed effect on the cell killing is summarized for each structure, and each lists the common name of the conjugate backbone. The addend function is listed for each.

**Extended Data Fig. 3.**
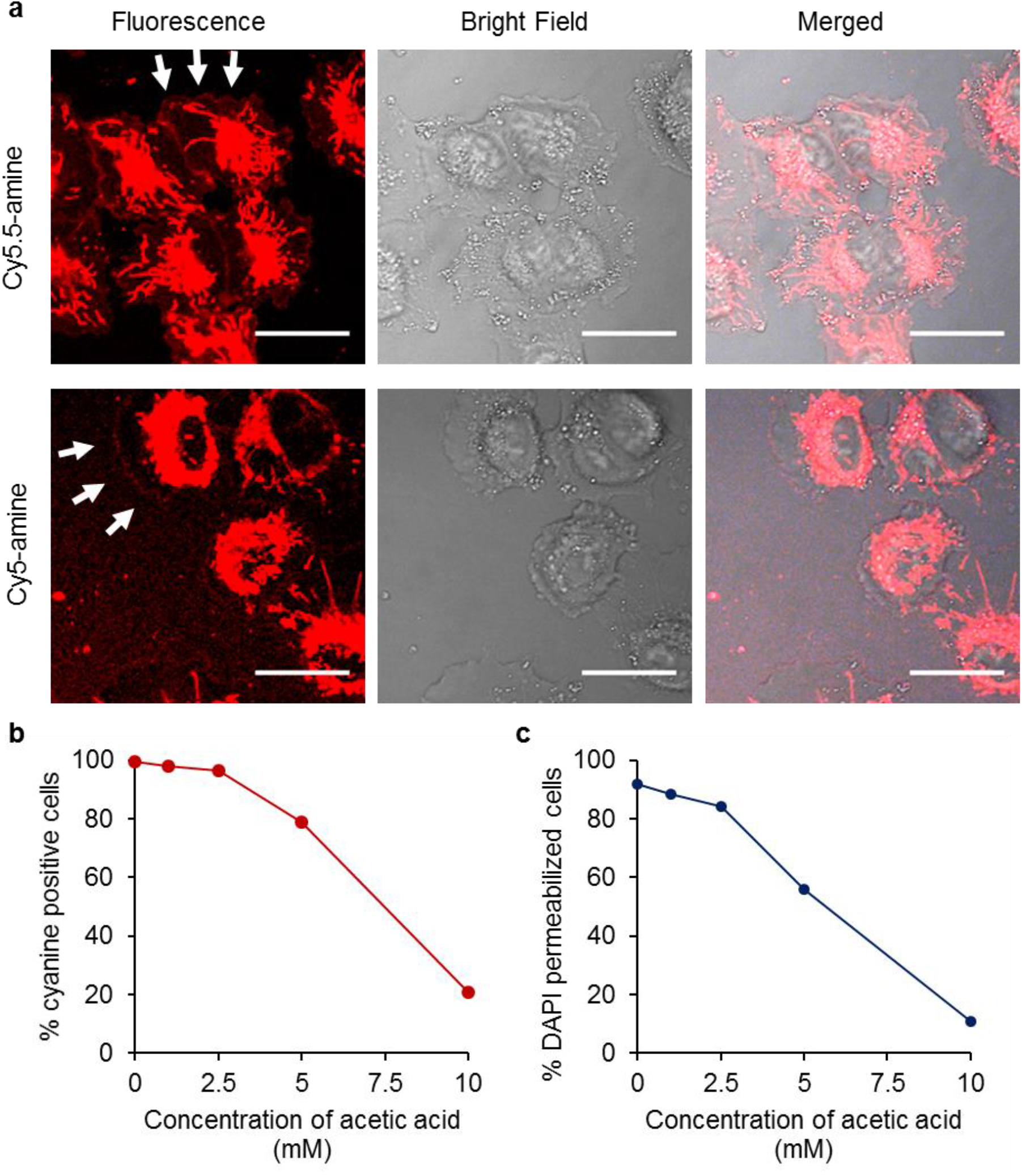
Binding of MJH into the external cellular membrane and into internal organelle membranes of A375 human melanoma cell line. (**a**) Fluorescence confocal microscopy imaging of MJH (Cy5.5-amine and Cy5-amine) loaded in A375 cells. C_loading_ = 2 μM, incubation time = 30 min, λ_ex_ = 640 nm, λ_em_ = 663-738 nm. The arrows are to indicate the position of the external cellular membrane and its staining with cyanine dyes (MJH). In total about 75 cells in average were analyzed in each condition in the confocal microscope in 5 different locations. Here representative images are shown. (**b**) Effect of the concentration of acetic acid in the binding of Cy7.5-amine to the A375 cells using flow cytometry analysis for quantification. Average of two experiments is shown (n = 2). (**c**) Effect of the concentration of acetic acid in the percentage of DAPI permeabilized cells using flow cytometry analysis for quantification. Cells were treated with 2 μM Cy7.5-amine, incubation for 30 min, then were illuminated with 730 nm light at 80 mWcm^-2^ for 60 s. Since acetic acid protonates the phosphates in the phospholipids, the Cy7.5-amine is not able to bind efficiently to lipid membranes. The lower binding of Cy7.5-amine to the cells is reflected in the lower permeabilization of the cells upon NIR light excitation. Average of two experiments is shown (n = 2) Scale bars = 25 μm.

**Extended Data Fig. 4.**
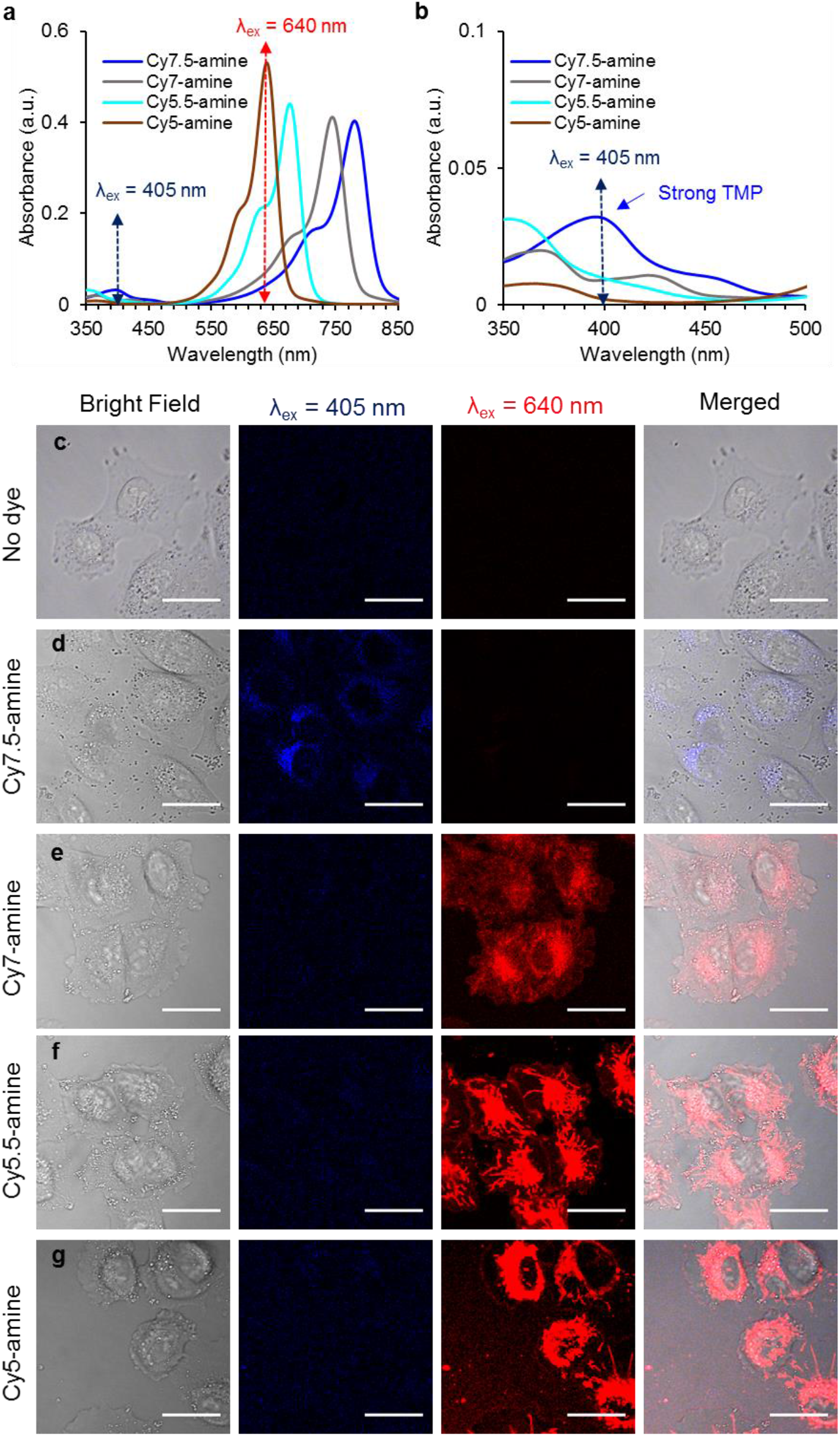
Absorption spectrum of MJH and confocal fluorescence microscopy of A375 cells in the presence of MJH. (**a**) Absorption spectrum of MJH showing the position of the excitation lasers that were used in the confocal microscope (λ_ex_ = 405 nm with λ_em_ = 425-475 nm or λ_ex_ = 640 nm with λ_em_ = 663-738 nm). (**b**) Expansion of the x and y axis of the absorption spectrum to observe the expected TMP (transversal molecular plasmon). The Cy7.5-amine shows a strong TMP (strong hybridization of longer C7 heptamethine bridge and larger benzoindole). Cy7-amine shows a weaker TMP and slightly shifted to ~375 nm because the C7 is hybridized to the smaller indole. Cy5.5-amine shows a strong TMP (larger benzoindole) but shifted to ~360 nm because of the hybridization with a weaker LMP (shorter C5 pentamethine). Cy5-amine shows little TMP because of the poor hybridization of the shorter C5 and smaller indole; this is the weakest combination because of the poor plasmonicity on both components. (**c**) Cells in the absence of dyes. (**d**) Cells in the presence of 2 μM Cy7.5-amine, 30 min of incubation. The emission of TMP mode can be observed at λ_ex_ = 405 nm in blue color. The excitation at 640 nm excites the tail of the LMP and produces a weak emission that is present but is difficult to see in the picture. (**e**) Cells in the presence of 2 μM Cy7-amine, 30 min of incubation. The emission from the LMP in Cy7-amine, since it is blue-shifted relative Cy7.5-amine, can be observed as red (λ_em_ = 663-738 nm). (**f**) Cells in the presence of 2 μM Cy5.5-amine, 30 min of incubation. The emission from the LMP is clearly visible in red. (**g**) Cells in the presence of 2 μM Cy5-amine, 30 min of incubation. The emission from LPM is clearly visible in red. The emission from TMP (at λ_ex_ = 405 nm) is not observed or is very weak in signal in **e-g** panel since the TMP is not present or shifted to other wavelengths on those molecules (Cy7, Cy5.5, and Cy5). In total about 75 cells in average were analyzed in each condition in the confocal microscope in 5 different locations. Here representative images are shown. Scale bars = 25 μm.

**Extended Data Fig. 5.**
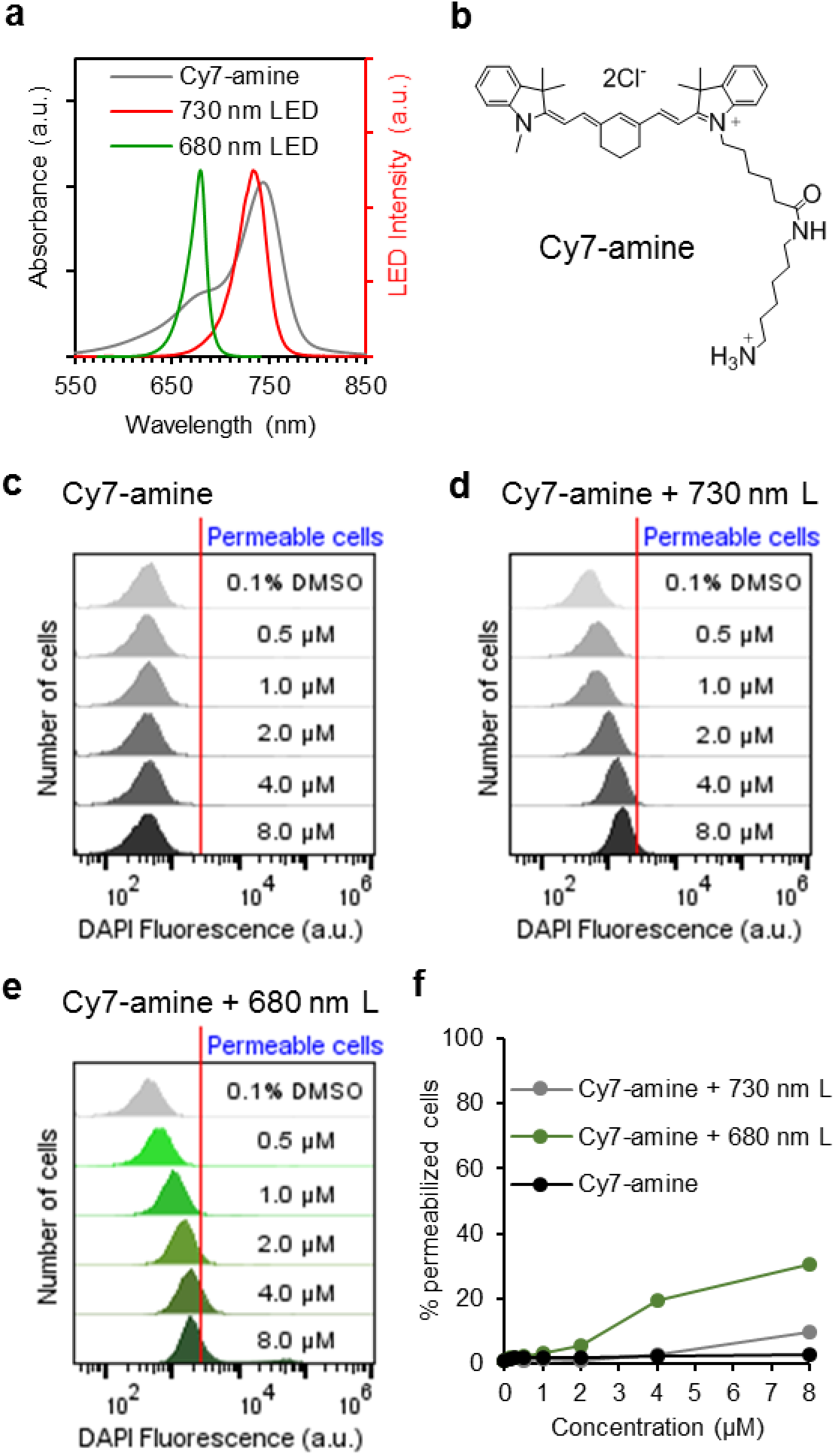
Excitation of the 680 nm vibronic shoulder in Cy7 improves the MJH effect for opening cell membranes in A375 cells. (**a**) Absorption spectra of Cy7-amine and overlaid with the spectral output of two LED lights: 730 nm and 680 nm. (**b**) Structure of Cy7-amine. (**c**) Flow cytometry analysis to measure the permeabilization of A375 cells in the presence of Cy7-amine without light. (**d**) Flow cytometry analysis to measure the permeabilization of A375 cells in the presence of Cy7-amine with 730 nm LED activation. (**e**) Flow cytometry analysis to measure the permeabilization of A375 cells in the presence of Cy7-amine with 680 nm LED activation. In all the flow cytometry analyses the red line represents the gating to discriminate between DAPI negative and positive cells (permeable). In all the cases the incubation with the cyanine was for 30 min and irradiation was with an equal light dose of 80 mWcm^-2^ for 10 min. The light dose was calibrated with an Optical Power Meter from Thorlabs, sensor model S302C and console model PM100D. (**f**) Percentage of permeabilized cells, the numbers are obtained from the flow cytometry. 10,000 cells are analyzed per each concentration.

**Extended Data Fig. 6.**
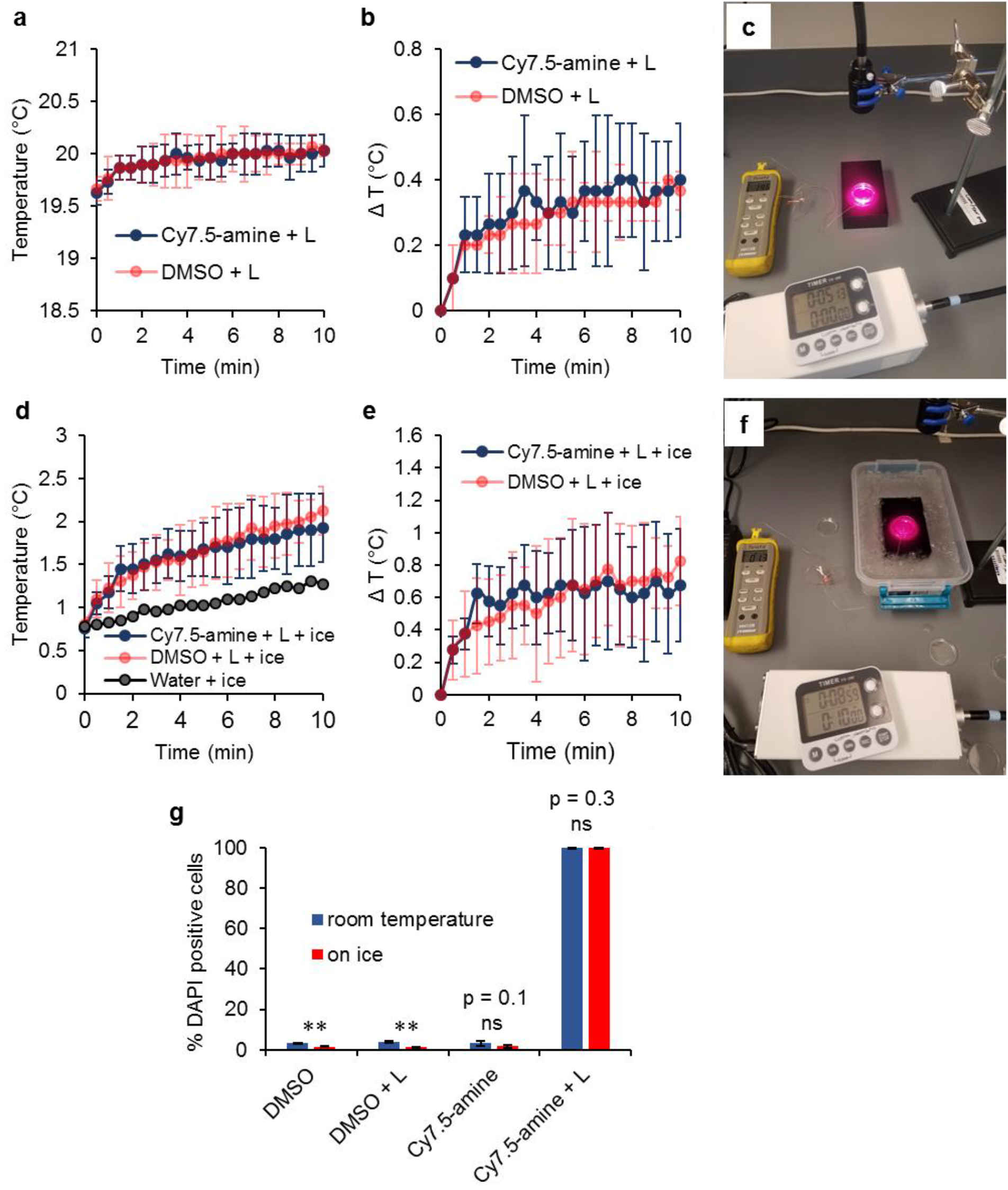
Temperature of the cell suspension while under light treatment. Temperature on the cell killing experiment using Cy7.5-amine. (**a-b**, **d-e**) No detection of heat production by Cy7.5-amine under NIR light treatment in the cell suspension (A375 cells) above the control. Temperature of the cell suspension (A375 cells) with 2 μM Cy7.5-amine and under illumination with 730 nm NIR light (80 mWcm^-2^ for 10 min). The temperature of the media was recorded when the experiment was done at room temperature (**a-c**) and when the cell suspension was placed in an ice bath (**d-f**). A picture of the experimental set up when done at room temperature is shown in **c** and in ice bath is shown in **f.** In **d** “water + ice” is the increase of the temperature because of the melting of ice without irradiation. In **e** the change of temperature is corrected by subtracting the temperature increase due to the melting of ice without illumination. The temperature of the cell suspension treated with NIR light and without Cy7.5-amine (DMSO + NIR light) correlates well with temperature profile in the suspension treated with NIR light containing 2 μM Cy7.5-amine (Cy7.5 + NIR light). There is no photothermal effect of Cy7.5-amine beyond the minimal heating caused by the light alone of ~0.5 °C. (**g**) Percentage of cells permeabilized to DAPI when the treatment was done at room temperature versus when was done placing the cell suspension on ice bath. In **a** and **b** 3 independent samples were processed and measured (n = 3). In **d** and **e** 4 independent samples were processed and measured (n = 4). In **g** 3 independent samples were processed and measured by flow cytometry (n = 3). In all the error bars are the standard deviation. *t*-test, two-tail, * p < 0.05, ** p < 0.01, *** p < 0.001 Statistical significance p < 0.05, ns = not significant.

**Extended Data Fig. 7.**
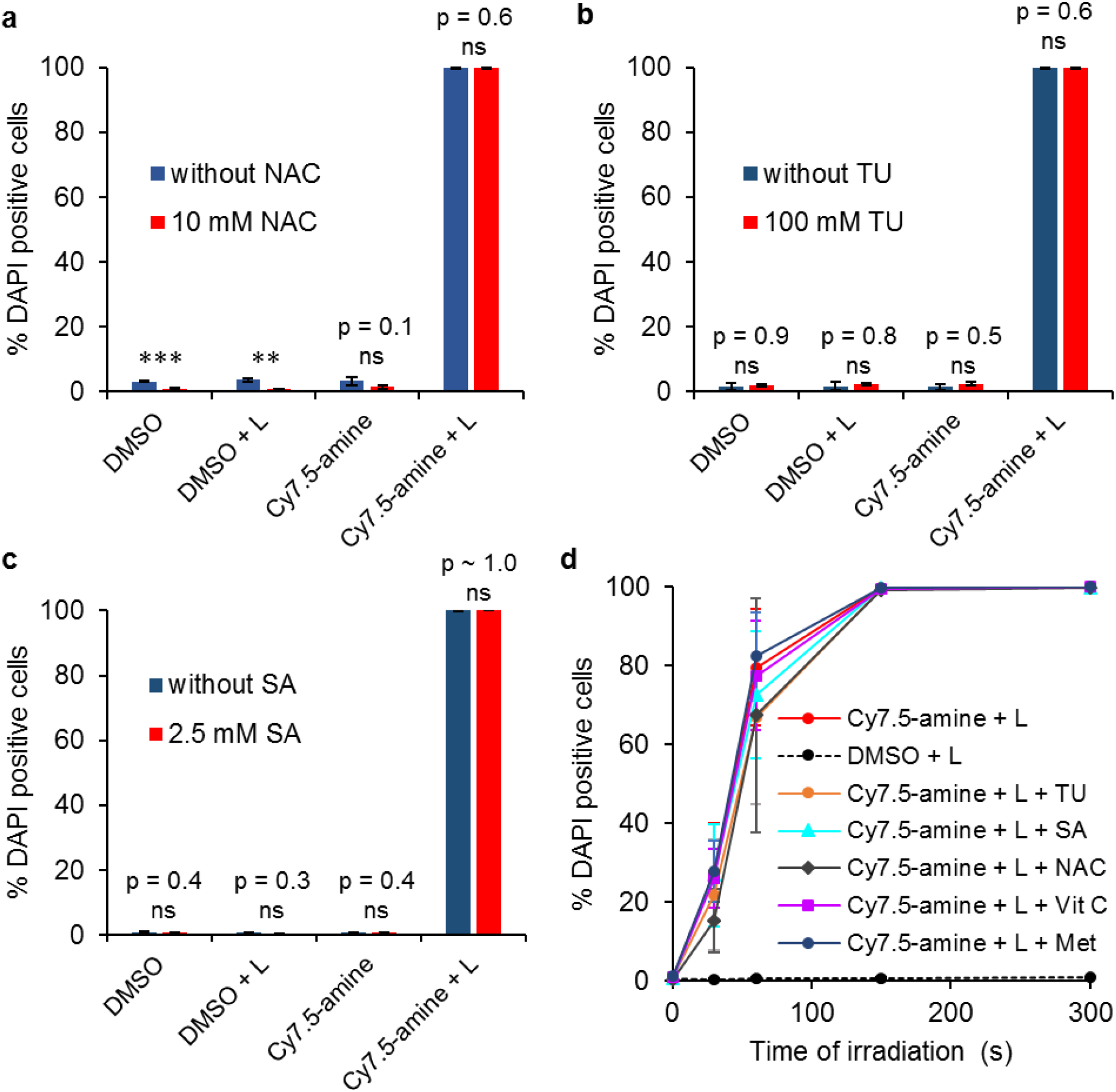
ROS effects on the cell killing using Cy7.5-amine. ROS scavengers do not retard the permeabilization of A375 cells to DAPI when treated with 2 μM Cy7.5-amine under illumination with 730 nm NIR light (80 mWcm^-2^ for 10 min). (**a**) Effect of 10 mM NAC (*N*-acetylcysteine). (**b**) Effect of 100 mM TU (thiourea). (**c**) Effect of 2.5 mM SA (sodium azide). (**d**) Effect of ROS scavengers at variable irradiation time of 730 nm NIR light at 80 mWcm^-2^. Five different scavengers were used: TU 100 mM, azide 2.5 mM, NAC 1 mM, Vit C (vitamin C) 5 mM and Met (methionine) 5 mM. DMSO control contains 0.1% DMSO in the media because DMSO is used to pre-solubilize the Cy7.5-amine stock solution at 2 mM and diluted to 1:1000 to obtain 2 μM Cy7.5-amine in media containing 0.1% DMSO. Experimental groups are: DMSO = 0.1% DMSO, DMSO+L = 0.1 % DMSO + NIR light treatment, Cy7.5 = 2μM Cy7.5-amine and Cy7.5+L = 2μM Cy7.5-amine + NIR light treatment. In all the plots 3 independent samples were processed and analyzed by flow cytometry (n = 3). In all the errors bars are the standard deviation. *t*-test, two-tail, * p < 0.05, ** p < 0.01, *** p < 0.001 Statistical significance p < 0.05, ns = not significant.

**Extended Data Fig. 8.**
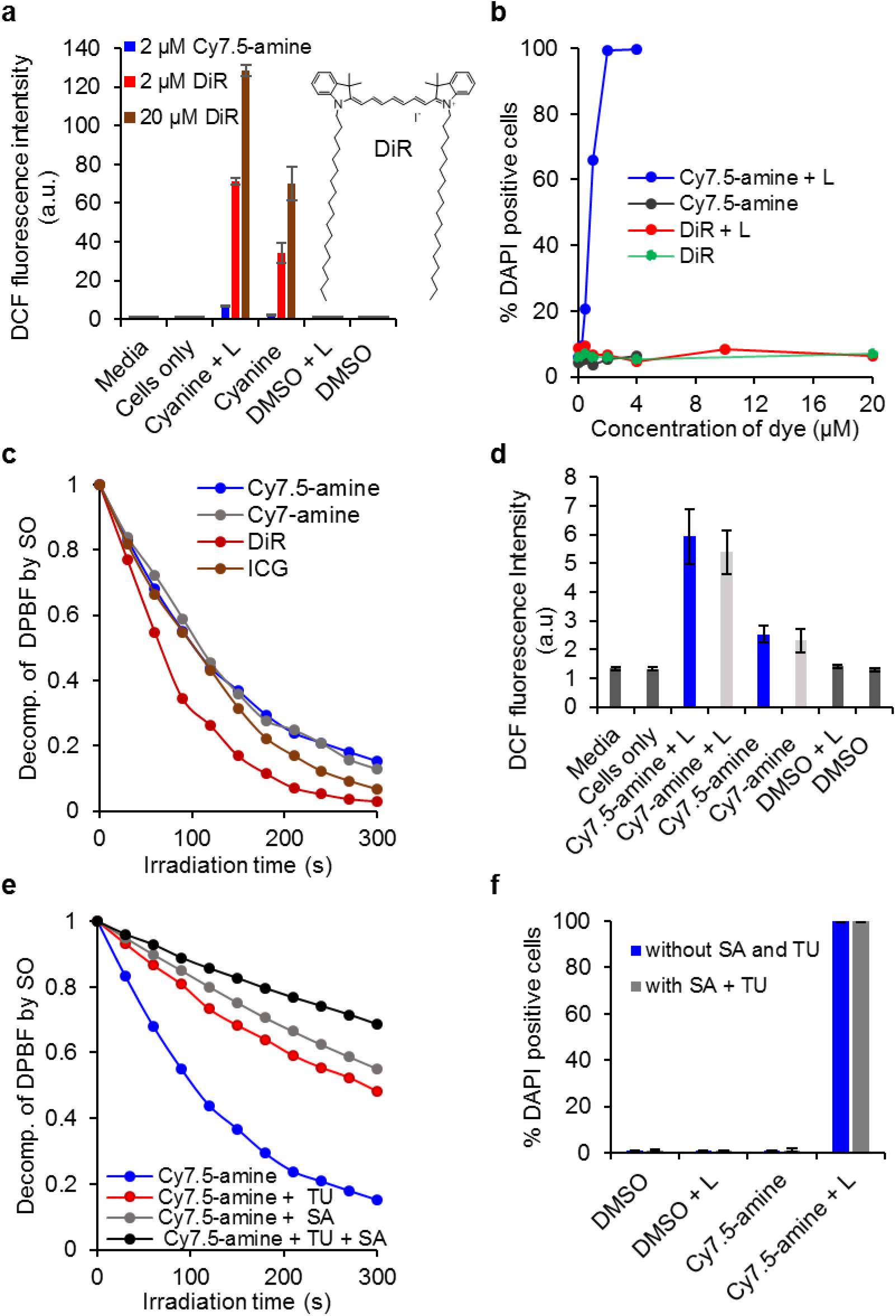
Quantification of ROS and singlet oxygen (SO) levels and their effect on the cell killing using Cy7.5-amine versus the cell-membrane-targeting DiR dye. (**a**) Measurement of ROS levels in A375 cell suspensions using 2’,7’-dichlorodihydrofluorescein diacetate as the ROS probe in the presence of 2 μM Cy7.5-amine versus 2 μM DiR and 20 μM DiR with and without light (L). (**b**) Percentage of DAPI positive cells as a function of the concentration quantified from the flow cytometry analysis. DiR that produced 10-20-fold more ROS than Cy7.5-amine did not permeabilize the A375 cells. This continues to support that the DiR structure is a weaker MJH with poorer VDA (See Extended Data Fig 2.) And more importantly, that ROS is not responsible for the permeabilization of DAPI into the cells. (**c**) Quantification of SO levels by the decomposition rate of DPBF (1,3-diphenylisobenzofuran) under light illumination in the presence of four cyanine dyes: 2.6 μM Cy7.5-amine, 2.6 μM Cy7-amine, 2.6 μM DiR and 2.6 μM ICG. DiR and ICG produces more SO than Cy7.5-amine or Cy7-amine yet DiR was unable to permeabilize the cells. (**d**) ROS levels in cells in the presence of Cy7.5-amine versus Cy7-amine. Cy7.5-amine and Cy7-amine produced approximately the same levels of SO and ROS yet Cy7.5-amine is a much stronger MJH in cell permeabilization (Fig. 1). (**e**) Shutting down the levels of SO generation by adding thiourea (TU = 100 mM) or sodium azide (SA = 2.5 mM) or a combination of TU 100 mM and SA 2.5 mM into the 2 μM Cy7.5-amine solution under LED illumination. (**f**) Effect of ROS scavenger combo (100 mM TU and 2.5 mM SA) on the percentage of DAPI positive cells as quantified from the flow cytometry analysis in the presence of 2 μM Cy7.5-amine with and without illumination: no difference observed in the cell permeabilization to DAPI. Unless otherwise specified, the light irradiations doses were 80 mWcm^-2^ for 10 min using a 730 nm LED. Except, in **b** the LED light (L) was a 740 nm light from Keber Applied Research Inc. at the same dose of 80 mWcm^-2^ for 10 min. This was done with a different LED because the data was collected in the early stage of the research, and we later moved to use the 730 nm LED for all the experiments. In panel **a**, the average is from 4 treated and measured samples (n = 4) for this the samples are quadruplicated in one 96 well. In panel **b** 5,000 cells are analyzed by flow cytometry for each concentration (one sample per each concentration n =1). In panel **c** the number of samples is n = 1. In panel **d**, 12 samples are processed and analyzed in each condition from 3 independent experiments, and in each experiment each condition is quadruplicated in a 96 well plate (n = 3 experiments x 4 samples per condition = 12). In panel **e** the number of samples n = 1. In panel **f**, n = 3 independent samples. In all the error bars are the standard deviations.

**Extended Data Fig. 9.**
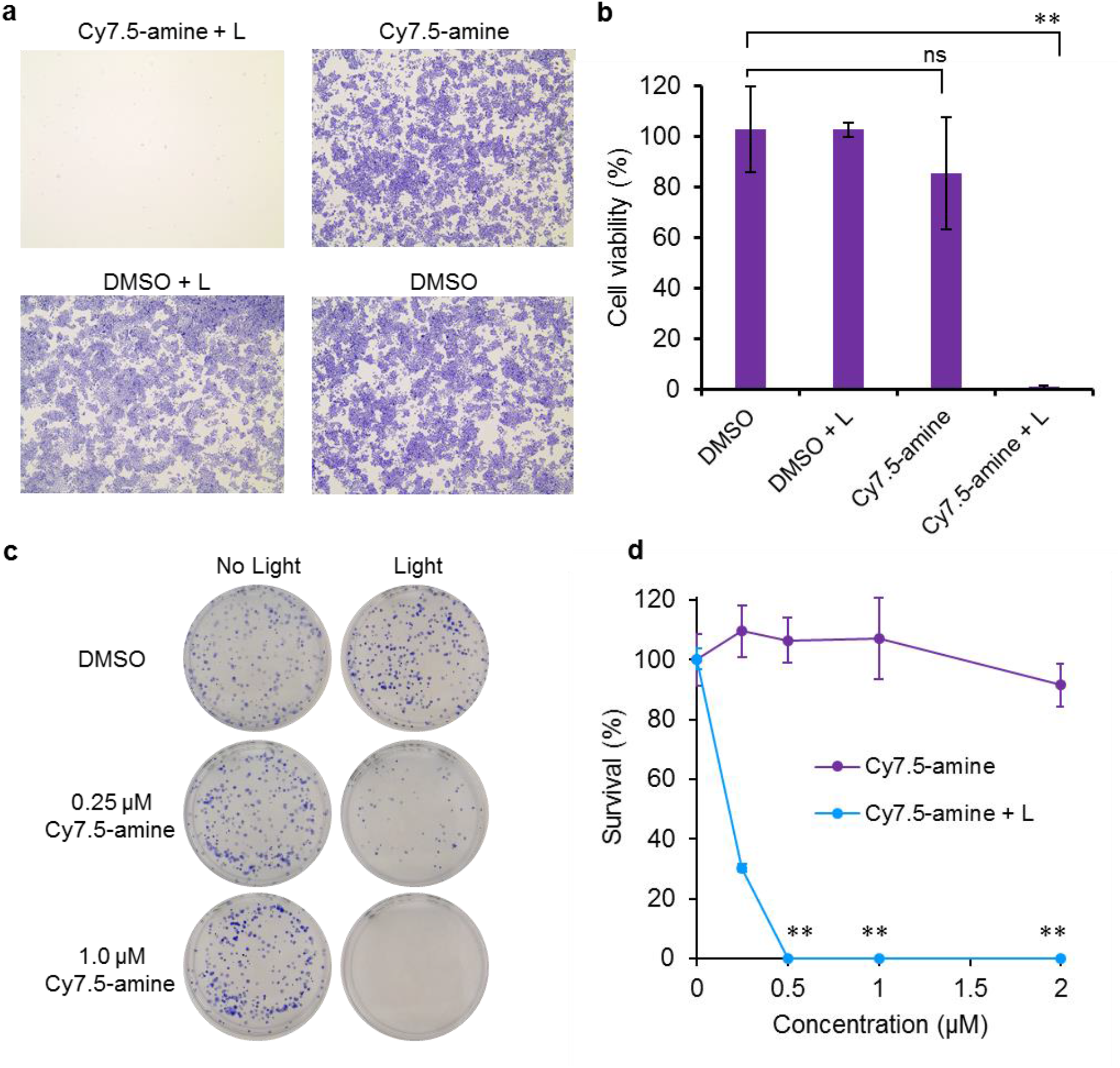
Quantification of cell death by crystal violet assay and clonogenic assay. A375 cells treated with Cy7.5-amine and 80 mWcm^-2^ of 730 nm NIR light for 10 min. (**a**) Representative microscopy picture of each condition in the crystal violet assay (n = 4). Experimental groups consist of: Cy7.5-amine + L = 2 μM Cy7.5-amine + 80 mWcm^-2^ of 730 nm NIR light for 10 min; Cy7.5-amine = 2 μM Cy7.5-amine; DMSO + L = 0.1% DMSO + 80 mWcm^-2^ of 730 nm NIR light for 10 min; and DMSO = 0.1% DMSO. (**b**) Crystal violet assay. Plot showing the quantification of the cell viability from the absorbance of crystal violet. Error bars are the standard deviations. Sample repetitions n = 4 for each condition in a 24 well plate (independent samples). (**c**) Clonogenic assay. Representative pictures showing the growth of cell colonies in the controls (0.1% DMSO with or without light) and complete eradication of A375 cells when treated with 1 μM Cy7.5-amine + light (80 mWcm^-2^ of 730 nm NIR light for 10 min). (**b**) Clonogenic assay. Quantification of the number cells forming colonies. The survival is the percentage of cells that formed colonies. Error bars are the standard deviation. The results are normalized relative to the DMSO control. Sample repetitions n = 3 (independent samples). *t*-test, two-tail, * p < 0.05, ** p < 0.01, *** p < 0.001 Statistical significance p < 0.05.

## References

1. García-López, V. et al. Molecular machines open cell membranes. Nature 548, 567–572 (2017).

2. Ayala-Orozco, C. et al. Visible-light-activated molecular nanomachines kill pancreatic cancer cells. ACS Appl. Mater. Interfaces 12, 410–417 (2020).

3. Guentner, M. et al. Sunlight-powered kHz rotation of a hemithioindigo-based molecular motor. Nat. Commun. 6, 8406 (2015).

4. Santos, A. L. et al. Hemithioindigo-based visible light-activated molecular machines kill bacteria by oxidative damage. Adv. Sci. 2203242 (2022).

5. Weissleder, R. A clearer vision for in vivo imaging. Nat. Biotechnol. 19, 316–317 (2001).

6. Liu, D. et al. Near-infrared light activates molecular nanomachines to drill into and kill cells. ACS Nano 13, 6813–6823 (2019).

7. Mustroph, H. & Towns, A. Fine structure in electronic spectra of cyanine dyes: are sub-bands largely determined by a dominant vibration or a collection of singly excited vibrations? ChemPhysChem 19, 1016–1023 (2018).

8. Cui, Y. et al. Molecular plasmon-phonon coupling. Nano Lett 16, 6390–6395 (2016).

9. Kong, F. F. et al. Probing intramolecular vibronic coupling through vibronic-state imaging. Nat. Commun. 12, 1280 (2021).

10. Chapkin, K. D. et al. Lifetime dynamics of plasmons in the few-atom limit. Proc. Natl. Acad. Sci. 115, 9134–9139 (2018).

11. Orlandi, G. & Siebrand, W. Theory of vibronic intensity borrowing. Comparison of Herzberg-Teller and Born-Oppenheimer coupling. J. Chem. Phys. 58, 4513 (2003).

12. Mishra, A., K. Behera, R., K. Behera, P., K. Mishra, B. & B. Behera, G. Cyanines during the 1990s: A Review. Chem. Rev. 100, 1973–2012 (2000).

13. Li, Y., Zhou, Y., Yue, X. & Dai, Z. Cyanine conjugates in cancer theranostics. Bioact. Mater. 6, 794–809 (2021).

14. Shi, C., Wu, J. B. & Pan, D. Review on near-infrared heptamethine cyanine dyes as theranostic agents for tumor imaging, targeting, and photodynamic therapy. J. Biomed. Opt. 21, 050901 (2016).

15. Lange, N., Szlasa, W., Saczko, J. & Chwiłkowska, A. Potential of cyanine derived dyes in photodynamic therapy. Pharmaceutics 13, 818 (2021).

16. Bilici, K., Cetin, S., Aydindogan, E., Yagci Acar, H. & Kolemen, S. Recent advances in cyanine-based phototherapy agents. Front. Chem. 9, 707876 (2021).

17. Štacková, L. et al. Deciphering the structure-property relations in substituted heptamethine cyanines. J. Org. Chem. 85, 9776–9790 (2020).

18. Malcioğu, O. B., Gebauer, R., Rocca, D. & Baroni, S. TurboTDDFT – A code for the simulation of molecular spectra using the Liouville–Lanczos approach to time-dependent density-functional perturbation theory. Comput. Phys. Commun. 182, 1744–1754 (2011).

19. Giannozzi, P. et al. QUANTUM ESPRESSO: a modular and open-source software project for quantum simulations of materials. J. Phys.: Condens. Matter 21, 395502 (2009).

20. Momma, K. & Izumi, F. VESTA 3 for three-dimensional visualization of crystal, volumetric and morphology data. J. Appl. Cryst. 44, 1272–1276 (2011).

