## Supporting Information for "Molecular Jackhammers Eradicate Cancer Cells by Vibronic-Driven Action"

**Cell binding competition of Cy5-amine against Cy7.5-amine.** The permeabilization activity of Cy7.5-amine in A375 cells was partially inhibited by the addition of Cy5-amine as shown in Fig. S1. In this test, Cy7.5-amine at 1 μM was used in the presence of 8 μM Cy5-amine.

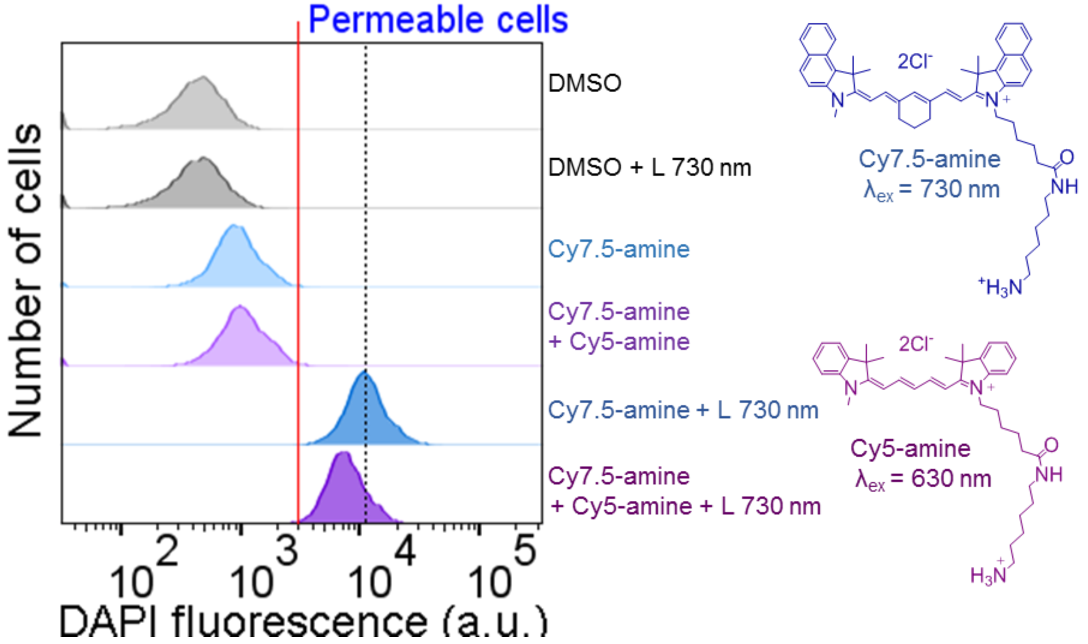


**Fig. S1 |** Flow cytometry analysis of Cy7.5-amine activity inhibition for permeabilization of cells using Cy5-amine as competitor molecule. Cy5-amine interacts with the cells and competes with Cy7.5-amine. The 730 nm LED excites Cy7.5-amine but does not excites Cy5-amine (the excitation of Cy5-amine using 730 nm LED is almost negligible). The concentration of Cy7.5-amine was 1 µM and Cy5-amine was 8 µM. The illumination was done with 730 nm LED, 80 mWcm$^{-2}$ for 10 min. The red line is to indicate the gate level for DAPI positive cells. The black dotted line is to guide the shifting position of the peak in presence of Cy5-amine. Three independent flow cytometry experiments were conducted (n = 3), a presentative is presented.

**Vibronic mode activation in ICG and effect on permeabilizing cancer cells.**

Another cyanine was also tried that has a vibronic absorption shoulder at 730 nm, indocyanine green (ICG), bearing alkylsulfonate addends, as shown in Fig. S2 a-b. The sulfonates will not interact with phospholipids in lipid bilayers. ICG is commonly used, and it is FDA-approved for intravenous injection. However, ICG was not able to permeabilize the cells under the same conditions in which Cy7.5-amine was tested, 2 μM and illumination for 10 min of 730 nm light at 80 mWcm$^{-2}$ (Fig. S2). The presence of 1% or less of DAPI positive cells is considered normal content of cell death in every cell batch and this is baseline in the control. Furthermore, ICG does not cause a photothermal effect at 2 μM since the temperature of the media remains at the baseline at 20 °C (Fig. S2 c). The photothermal effects of ICG were studied at the high concentration range of 8 μM up to 400 μM (Fig. S2 c). Below 32 μM, from 0 to 16 μM, there was no observed cell permeabilization to DAPI with or without the light treatment (Fig. S2 d, f). In the range of 32 μM up to 400 μM, there was observed permeabilization of the cells to DAPI from 2% up to 14%, respectively (Fig. S2 e, f). This cell permeabilization was caused by the photothermal effect since the medium showed a significant temperature increase in this range of concentrations. The permeabilization of the cells by the photothermal effect with ICG reached its maximum at 200 μM because of the attenuation of the light by ICG; the transmittance of the light is compromised as the ICG concentration increases.

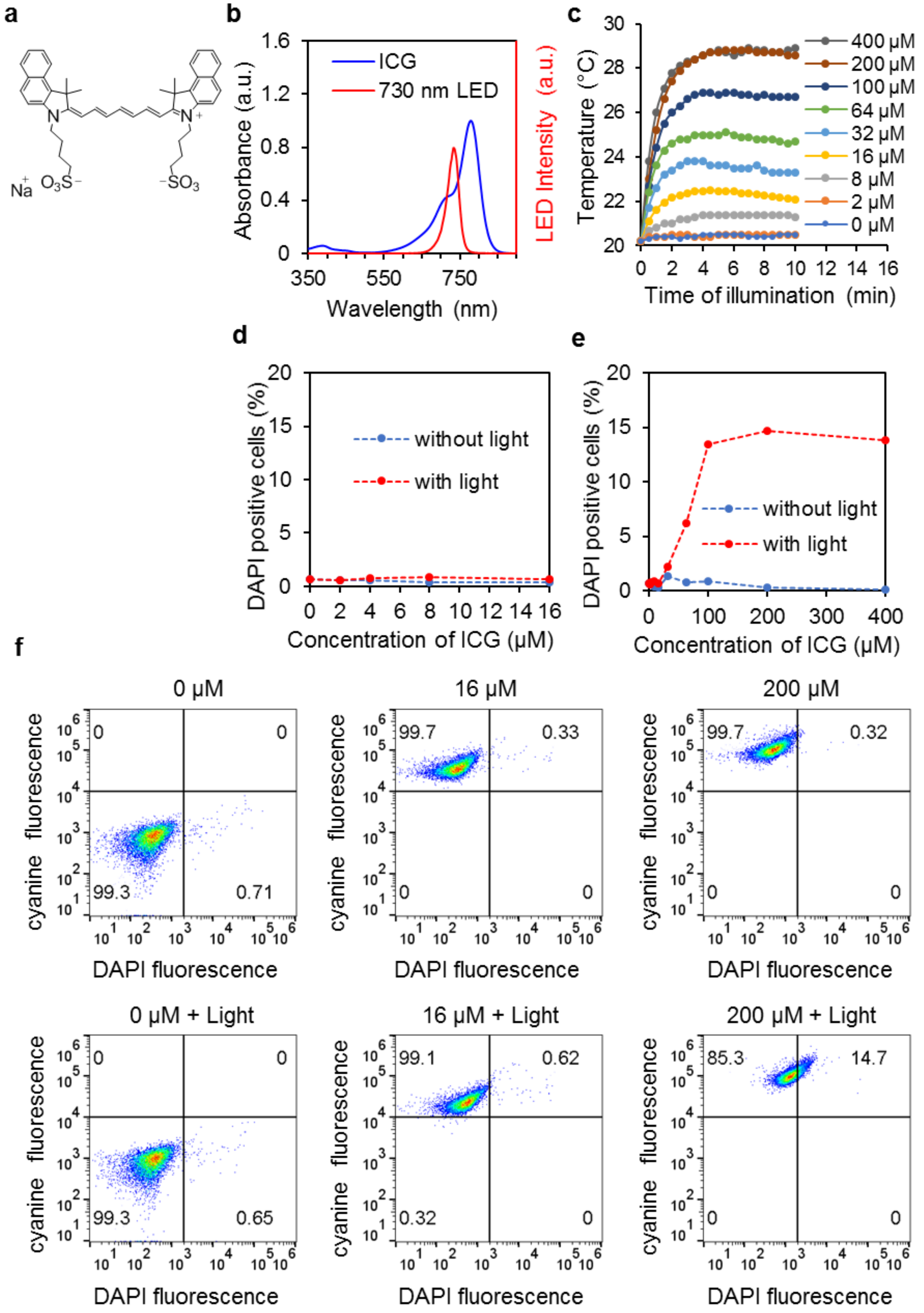
a
b
ICG
730 nm LED
Absorbance (a.u.)
LED Intensity (a.u.)
Wavelength (nm)
c
400 μM
200 μM
100 μM
64 μM
32 μM
16 μM
8 μM
2 μM
0 μM
Temperature (°C)
Time of illumination (min)
d
without light
with light
DAPI positive cells (%)
Concentration of ICG (μM)
e
without light
with light
DAPI positive cells (%)
Concentration of ICG (μM)
f
0 μM
16 μM
200 μM
0 μM + Light
16 μM + Light
200 μM + Light
cyanine fluorescence
DAPI fluorescence

**Fig. S2 | Effect of indocyanine green (ICG) activated by 730 nm light on A375 melanoma cells.** (**a**) Structure of ICG. (**b**) Absorption spectrum of ICG in water with 0.1% DMSO. The LED spectrum is also shown. (**c**) Photothermal heating effect of ICG at various concentrations and illuminated by 730 nm light at 80 $mWcm^{-2}$ over time from 0 to 10 min. (**d**) Percentage of permeabilized cells (DAPI positive cells) in the range of ICG concentration from 0 to 16 μM. Cells were illuminated with 730 nm light at 80 $mWcm^{-2}$ for 10 min. (**e**) Percentage of permeabilized cells (DAPI positive cells) in the range of ICG concentration from 0 to 400 μM. Cells were illuminated with 730 nm light at 80 $mWcm^{-2}$ for 10 min. (**f**) Representative flow cytometry plots showing the DAPI positive gates from where **d** and **e** were constructed. The numbers inside the gates (four quadrants) in the flow cytometry plot represent the percentage of cells in each gate: ICG and DAPI negative (left bottom), ICG positive and DAPI negative (left top), ICG negative and DAPI positive (right bottom), and ICG positive and DAPI positive (top right). 10,000 cells are analyzed in each concentration. One sample is treated and analyzed for each concentration.

**Flow cytometry data processing and gating strategy.** The flow cytometry data was analyzed using FlowJo software version 10.5.3. The cells were plot using the forward scattering area (FSC-A) versus side scattering area (SSC-A) as shown in Fig. S3 a. Then, a wide polygonal gate was drawn as shown in Fig. S3 a since we were interested to detect any cell death after treatment, we were not only interested to measure healthy cell populations. Sometimes, we observed that some death cell population upon treatment could shift outside the gate if a narrow gate was used. Then, the singlet cells were selected by plotting the FSC-A versus the forward scattering height (FSC-H) as shown in Fig. S3 b. Then, the singlets were selected and the DAPI fluorescence intensity was plotted versus the SSC-A and the gate for DAPI positive cells was drawn as shown in Fig. S3 c and the analysis was applied to all samples in the group. In Fig. S3 d can be observed a DAPI positive population in the same analysis group. This analysis was applied to calculate the percentage of DAPI positive cells.

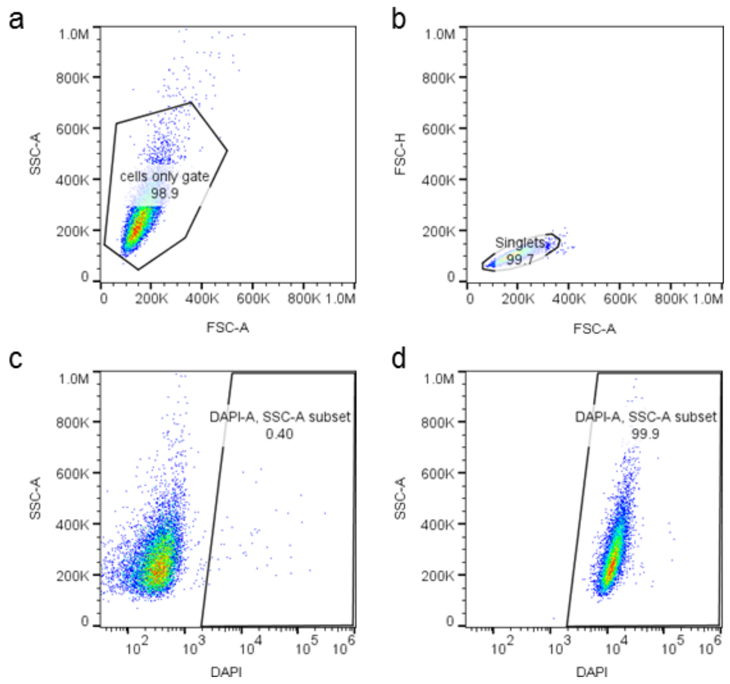


**Fig S3 | Flow cytometry data processing and gating strategy.** Data analysis was done using FlowJo version 10.5.3. a) Selection of the cell population by plotting forward scattering area (FSC-A) versus side scattering area (SSC-A). b) Selection of single cells by plotting FSC-A versus FSC-height (FSC-H). c) Gating of DAPI positive cells in a control sample containing 0.1% DMSO. d) Application of the same gate conditions as shown in c to a DAPI positive sample which was treated with Cy7.5-amine and 730 nm light. The number in the inset in each gate shows the percentage of positive cells.
